# Aptamer-Targeted PLGA Nanoparticles for Biofilm-Directed Intra-articular Antibiotic Delivery in Prosthetic Joint Infection

**DOI:** 10.64898/2026.10.23.760933

**Authors:** Zahra Eskandari, Yingfang Fan, Keita Fujino, Amita Sekar, Philip Ezekiel Tan, Fawaz Ben Malick, Yiwei Yan, Mehmet D. Asik, Orhun Muratoglu, Ebru Oral

## Abstract

**Introduction:** Prosthetic joint infection (PJI) is a severe complication of orthopedic surgery, driven by biofilm-forming bacteria such as *Staphylococcus aureus*. Conventional antibiotics are limited in penetrating biofilms or remaining localized at the infection site.

We engineered a targeted nanocarrier system by conjugating DNA aptamers specific to *S. aureus* biofilm components onto poly(lactic-co-glycolic acid) (PLGA) nanoparticles (NPs) encapsulating vancomycin. PLGA NPs were synthesized using flow-focusing microfluidics, followed by surface functionalization via EDC/NHS chemistry. Aptamer conjugation was validated using gel electrophoresis and DLS. Targeting efficiency and antimicrobial activity were evaluated through fluorescence microscopy, bacterial viability assays and *in vitro* biofilm models. Biodistribution and joint retention were assessed using fluorescent aptamer-conjugated NPs following intra-articular injection in rat knee joints.

**Results:** The optimized vancomycin-loaded NPs exhibited uniform morphology, high vancomycin loading capacity (7.9 ± 2.2%), and a sustained biphasic release profile with an early high-release phase followed by slower extended release. PK modeling indicated prolonged intra-articular drug retention with concentrations maintained above the minimum inhibitory concentration (MIC) for an extended period. Aptamer conjugation preserved nanoparticle stability and enhanced binding and retention within *S. aureus* biofilms *in vitro*, supporting improved biofilm localization. In-vivo fluorescence imaging demonstrated high local retention of aptamer-conjugated NPs, supporting localized joint retention with limited off-target tissue signal.

**Conclusions:** *S. aureus*-specific aptamer conjugated nanoparticles (SA Apt-P(V)) represent a targeted intra-articular delivery platform with potential for localized PJI therapy. The combined in-vitro and in-vivo findings highlight their potential to improve local therapeutic concentration, minimize systemic exposure, and enhance infection control within prosthetic joints.

## 1. Introduction

Prosthetic joint infection (PJI) remains one of the most serious and costly complications of arthroplasty. Contemporary epidemiology shows that PJI occurs in 1–2% of primary hip and knee replacements and at higher rates after revision surgery, [1, 2]. PJI is a major driver of reoperations: registry and observational studies report that infection accounts for ∼25–30% of revision total knee arthroplasties and a substantial proportion of hip revisions [3, 4]. Together, these numbers underscore the clinical and economic urgency of improving early detection and targeted treatment strategies for PJI. PJI management is complicated by the ability of pathogens, particularly *Staphylococcus aureus (S. aureus),* to form biofilms on implant surfaces. These biofilms protect bacteria from host immune responses and markedly reduce antibiotic susceptibility, making eradication extremely difficult [5, 6].

Current treatments rely primarily on systemic antibiotic administration, often combined with surgical intervention [7]. Procedures with low invasiveness such as debridement, antibiotics, and implant retention (DAIR) or polyethylene liner exchange aim to preserve the existing prosthesis. However, these approaches have limited long-term success [8] particularly in the presence of biofilm. The gold-standard two-stage revision involves removal of the infected implant, aggressive debridement, prolonged systemic antibiotic therapy, and eventual reimplantation. While the most effective (70-85% success rate), it is highly invasive, costly, and physically demanding on patients [4, 9, 10]. Given the morbidity and recurrence rates associated with existing methods, there is an urgent need for adjunct or alternative non-invasive therapies that can disrupt biofilms and enhance antibiotic efficacy.

Antibiotic therapy remains central to PJI management, with systemic agents like ciprofloxacin and fifth-generation cephalosporins used to target a broad bacterial spectrum. Staphylococcal species—particularly methicillin-sensitive and methicillin-resistant *S. aureus* (MSSA and MRSA)—are the leading causes of PJI, with MRSA infections resulting in worse clinical outcomes [11]. Vancomycin is a preferred antibiotic for treating infections caused by methicillin-resistant *S. aureus* (MRSA), one of the primary etiological agents of PJI [12]. However, its efficacy is limited in biofilm-rich environments due to poor penetration into the dense extracellular matrix and reduced activity against sessile bacteria. Moreover, systemic vancomycin administration is constrained by nephrotoxicity, the need for therapeutic drug monitoring, and limited bioavailability at the infected joint space [13, 14]. Consequently, there has been growing interest in localized drug delivery systems that can provide sustained, targeted antibiotic release directly at the site of infection.

Poly(lactic-co-glycolic acid) (PLGA), a biodegradable and biocompatible copolymer, has been widely used in controlled-release drug delivery systems, including microparticles, gels, and implants [15, 16]. Its clinical safety and efficacy have been demonstrated by several FDA-approved PLGA-based products [15, 16]. Several studies have successfully demonstrated the encapsulation of vancomycin into PLGA-based systems for sustained release and improved therapeutic profiles [17–20]. Among these, a previous study developed PLGA microparticles that served as effective local depots for vancomycin delivery in joint infections [19]. However, due to their relatively large size, these microparticles lack the ability to efficiently penetrate bacterial biofilms.

In contrast, PLGA nanoparticles provide additional advantages, particularly in the context of biofilm-associated infections. Their smaller size facilitates deeper diffusion into biofilms and enhanced interaction with bacterial cells embedded in the extracellular matrix [21, 22]. Furthermore, their higher surface-area-to-volume ratio enables more efficient surface functionalization and drug loading. PLGA nanoparticles loaded with vancomycin have been shown to maintain sustained drug release profiles while improving the therapeutic efficacy against *S. aureus* biofilms both *in vitro* and *in vivo* [20, 23–26]. In another study, PLGA nanoparticles enhanced vancomycin delivery to MRSA biofilms, resulting in improved bacterial killing compared to free drug [20].

To further augment the specificity of nanoparticle-based systems, aptamers have emerged as highly promising targeting ligands. Aptamers are short, single-stranded DNA or RNA oligonucleotides that can bind to specific molecular targets with high affinity and specificity, comparable to antibodies [27, 28]. Unlike antibodies, aptamers are synthetically produced, exhibit lower immunogenicity, and can be chemically modified for enhanced stability and conjugation to delivery systems [28]. Aptamer-conjugated nanoparticles have been studied in various therapeutic contexts, including cancer, thrombosis, and viral infections [29, 30]. Their application in antimicrobial therapy is relatively recent but rapidly expanding, with studies demonstrating their ability to direct nanocarriers specifically to bacterial cells or biofilm components [31, 32].

Despite the increasing interest, there is limited research on the use of aptamer-conjugated PLGA nanoparticles for targeted treatment of PJI, especially with *S. aureus* biofilms. In a recent study, Ommen et al. developed a liposomal system functionalized with a biofilm-specific aptamer for *S. aureus* and showed enhanced accumulation within biofilms and increased bacterial killing *in vitro* [32]. These findings suggest that aptamer-mediated targeting can significantly improve nanoparticle localization and drug delivery to biofilm-infected tissues. In this study, we developed vancomycin-loaded PLGA nanoparticles conjugated with a biofilm-targeting aptamer specific to *S. aureus* (SA Apt-P(V)). Our aim was to improve therapeutic delivery by combining nanoparticle-based controlled release, nanoscale biofilm interaction, and the molecular specificity offered by aptamers. This strategy was evaluated through in vitro studies of antimicrobial activity and biofilm targeting and in vivo studies of joint retention, biodistribution, and biocompatibility in a rat model of joint infection. By leveraging the enhanced permeability of nanoparticles and the binding specificity of aptamers, we propose a novel and clinically translatable nanotherapeutic platform for treating S. aureus-associated PJIs.

## 2. Methods

Poly(lactic-co-glycolic acid) (PLGA; Resomer^®^ RG 503H, lactide:glycolide ratio 50:50, acid-terminated, inherent viscosity 0.32–0.44□dL/g) was obtained from Evonik Industries (Germany). Vancomycin hydrochloride was obtained from Hospira, Inc. (USA). Sodium chloride, ethyl acetate, chloroform, Rhodamine B base (≥97% dye content) and Polyvinyl alcohol (PVA; Mw 13,000–23,000, 87–89% hydrolyzed) were purchased from Sigma-Aldrich (USA). The carbodiimide coupling agents 1-ethyl-3-(3-dimethylaminopropyl)carbodiimide hydrochloride (EDC) and N-hydroxysuccinimide (NHS), UltraPure™ DNase/RNase-free distilled water and phosphate-buffered saline (PBS) were purchased from Thermo Fisher Scientific (USA). The Cy5.5 amine and Cy5.5 carboxyl dyes were obtained from BroadPharm (USA).

Custom single-stranded DNA aptamers were synthesized by Integrated DNA Technologies (IDT, USA). The biofilm-targeting aptamer (SA20) and its Cy5- or Cy5.5-modified versions were used. Its sequence was: (GCAATGGTACGGTACTTCCGCGCCCTCTCACGTGGCACTCAGAGTGCCGGAAGTTCTGC GTTATCAAAAGTGCACGCTACTTTGCTAA). A scrambled-sequence control aptamer with identical length, ratio of nucleotides and modifications was used. Its sequence was: (AGTTTCAGGCCGGCCATCAAGTCGCCCAGACCAGCGAGGCATTACATGGCTTCATCTTTACTTATGGCCCGGAGTTGGCTTAGAATCC).

### 2.1 Development of Nanoparticles

Vancomycin-loaded poly(lactic-co-glycolic acid) (PLGA) nanoparticles (P(V) NPs) were synthesized using a water-in-oil-in-water (W/O/W) double emulsion technique in a flow focusing microfluidic system (Supplementary Fig. 1). The primary aqueous phase (W1) consisted of 0.6□mL vancomycin (100□mg/mL in water), while the organic phase (O) contained 2.4□mL PLGA (15□mg/mL in ethyl acetate). The secondary aqueous phase (W2) was composed of 2% polyvinyl alcohol (PVA) with 0.5□M NaCl in water. Set flow rates were 0.2□mL/min (W1), 0.8□mL/min (O), and 20□mL/min (W2) for maximized nanoparticle formation. The emulsion was kept under a chemical hood on a shaker at 250□rpm until solvent evaporation was complete.

After overnight solvent evaporation, nanoparticles were collected by centrifugation at 20,000 RCF at 12□°C for 15 minutes. The supernatant was discarded, and the pellet was washed with 10□mL PBS, resuspended, and centrifuged again under the same conditions. This washing process was repeated three times to remove residual surfactants and unencapsulated drug. The final nanoparticle pellet was resuspended in 10□mL DI water, frozen overnight at −30□°C, and lyophilized using a lyophilizer (BenchTop Pro with Omnitronics™, SP Scientific, USA) for 48 hours until complete drying was achieved. The lyophilized nanoparticles were stored at −30□°C until further use.

Blank PLGA NPs for degradability studies were prepared using the same microfluidic double emulsion method described above, without the inclusion of vancomycin.

Blank NPs used for biodistribution and Rhodamine B loaded PLGA (P(RhB)) used for imaging studies, were prepared via nanoprecipitation using the NanoAssemblr^®^ Benchtop system (Precision NanoSystems, Canada). A NanoAssemblr™ Ignite cartridge was used. The formulation consisted of PLGA (100 mg) and RhB (0.5□mg) in ethyl acetate as the organic phase, and 2% PVA in water as the aqueous phase. The organic and aqueous phases were introduced at flow rates of 1□mL/min and 3□mL/min, respectively, under continuous flow conditions. For clarity, all nanoparticle formulations and abbreviations used in this study are summarized in Table 1.

**Table 1.** Summary of nanoparticle formulations and abbreviations used throughout the manuscript.

| **Formulation full name** | **Abbreviation** | **Purpose in study** |
| --- | --- | --- |
| **Vancomycin loaded PLGA nanoparticles** | P(V) | Antibiotic delivery and release studies |
| **SA specific Aptamer-conjugated vancomycin loaded PLGA nanoparticles** | SA Apt-P(V) | Targeted biofilm binding |
| **Rhodamine B loaded PLGA nanoparticles** | P(RhB) | In vitro biofilm retention visualization |
| **Aptamer-conjugated RhB loaded PLGA nanoparticles** | SA Apt-P(RhB) | Targeted biofilm retention imaging |
| **Cy5-labeled SA Aptamer-conjugated blank PLGA nanoparticles** | Cy5-SA Apt-NP | Fluorescence microscopy biofilm targeting |
| **Cy5-labeled Scr Aptamer-conjugated blank PLGA nanoparticles** | Cy5-Scr Apt-NP | Fluorescence microscopy biofilm targeting |
| **Cy5.5-labeled SA Aptamer blank PLGA nanoparticles** | Cy5.5-SA Apt-NP | In vivo retention in infection model |
| **Cy5.5-labeled scrambled-aptamer blank PLGA nanoparticles** | Cy5.5-Scr Apt-NP | In vivo control |

### 2.2 Nanoparticle Characterization

Nanoparticle size and surface charge were measured using a Zetasizer (Malvern ZSP) at a detection angle of 173° at a nanoparticle concentration of 0.5 mg/mL in distilled water. Morphology analysis was conducted by mounting lyophilized nanoparticles on carbon tape affixed to aluminum stubs, followed by sputter-coating with a 5 nm platinum/palladium (Pt/Pd) layer, and imaging under high vacuum using a Gemini 560 Scanning Electron Microscope (SEM).

### 2.3 Loading Capacity (LC%) Determination

To determine vancomycin loading capacity (LC%), 5–10 mg of lyophilized nanoparticles were precisely weighed and dissolved in 2 ml chloroform. The mixture was vortexed for 30 seconds, sonicated for 3 minutes, and stirred with a magnetic stirrer for 30 minutes to ensure complete dissolution. Next, 5 mL of water was added, followed by 30 seconds of vortex mixing and 2 hours of stirring. Then, the stirrer was then turned off, and the sample was left undisturbed for 30 minutes until full phase separation was achieved. A 1 mL aliquot of the aqueous phase was collected, centrifuged at 20,000 RCF for 5 minutes, and the supernatant was analyzed in 5 replicates using a UV-Vis Synergy H1 plate reader (BioTek, USA) at 280 nm to quantify vancomycin concentration. The loading capacity (LC%) was calculated by dividing the detected vancomycin amount by the initial nanoparticle mass.

### 2.4 Drug Release

Vancomycin release from P(V) NPs was evaluated at 37°C with complete media replacement at each time point. Nanoparticles (45 mg in 2 mL) were dispersed in distilled water and incubated. At each timepoint, the entire external aqueous phase was removed and replaced with fresh distilled water to maintain sink conditions. Drug release was quantified by measuring absorbance at 280 nm using a (Synergy H1, BioTek, USA) UV-Vis plate reader, and concentrations were determined based on a vancomycin calibration curve. All experiments were performed in triplicate. To define the low-concentration analytical range for terminal release samples, a low-range vancomycin UV–Vis calibration was performed in distilled water, matching the release medium. The limit of detection (LOD) and limit of quantification (LOQ) were calculated as 3.3σ/S and 10σ/S, respectively, where σ is the standard deviation of the blank and S is the calibration slope.

### 2.5 Pharmacokinetic Modeling of Intra-articular Exposure

Intra-articular vancomycin concentrations were estimated using a compartmental pharmacokinetic model based on the experimentally measured cumulative release profile *in vitro*. Release data were scaled to a total administered polymer mass of 160 mg PLGA NPs. A one-compartment joint space model with first-order elimination was assumed, using a joint volume of 2 mL and an elimination half-life (t_½_) of 6 h, consistent with prior intra-articular clearance estimates. Modeled concentration–time profiles were generated by convolving the release rate with the elimination function, and pharmacokinetic parameters (C_max_, T_max_, and AUC) were calculated from the simulated profiles.

### 2.6 PLGA Degradation

To evaluate PLGA nanoparticle degradation, the pH of the release media collected during the release studies was monitored over time using a SevenMulti pH meter (Mettler Toledo, Switzerland) equipped with an InLab Expert Pro electrode, at a constant temperature of 25 °C. To isolate the contribution of the polymer, blank PLGA NPs (no vancomycin) were prepared at the same nanoparticle concentration and incubated under identical conditions as vancomycin-loaded formulations. At each sampling time point, after media replacement, the pH of the release medium was recorded for both blank and loaded nanoparticles to generate comparative pH–time profiles.

### 2.7 Aptamer Conjugation to PLGA NPs and Purification

Aptamer-conjugated PLGA NPs were prepared using EDC/NHS chemistry to activate the carboxyl groups on the nanoparticle surface. For every 10 mg/mL of nanoparticles, 400 mM EDC and 480 mM NHS were used. Before conjugation, nanoparticles were redispersed in UltraPure™ DNase/RNase-free distilled water for 5 minutes. EDC (400 mM) was added first, followed by NHS (480 mM) after 10 minutes, and the reaction was allowed to proceed for 20 minutes at room temperature under continuous stirring at 700 rpm on a magnetic stirrer.

Following activation, the reaction mixture was filtered and purified using Amicon® Ultra Centrifugal Filters (100 kDa MWCO) at 7,500 rpm for 15 minutes. The nanoparticles were then washed with 1 mL of DNAse/RNAse-free water, and the washing and centrifugation process was repeated three additional times under the same conditions. After the second washing step, eluates were collected for quantification of aptamer attachment and evaluation of washing efficiency. The final nanoparticle suspension was collected in the same volume of DNase/RNase-free distilled water as at the start of the reaction.

Parallel to this process, aptamers were folded before conjugation. The aptamer stock solution containing 5 mM MgCl_2_ was heated at 95°C for 5 minutes, followed by rapid cooling on ice for 15 minutes to promote proper folding. The folded aptamers were then added to the activated nanoparticles at a 1:10 molar ratio of aptamer to initial nanoparticles, and the reaction mixture was stirred at room temperature for 2 hours (700 rpm on magnetic stirrer) to facilitate aptamer binding.

Following incubation, the PLGA NP-aptamer bioconjugates were washed again using Amicon ultracentrifugation under the same conditions as before. Filtrates from these wash steps were collected to assess the extent of aptamer attachment and washing efficiency. The final aptamer-conjugated nanoparticles were collected in 5 mM MgCl_2_ and stored at 4°C until further use.

The same attachment method with a 1:30 molar ratio was used to prepare fluorescently labeled formulations: Cy5-tagged aptamer NPs (Cy5-SA and Cy5-Scr) for *in vitro* studies and Cy5.5– tagged counterparts (Cy5.5-SA and Cy5.5-Scr) for *in vivo* imaging. For simplicity, these formulations are referred to as fluorescent NPs throughout the manuscript.

### 2.8 Attachment Efficiency and Washing Validation

To evaluate the efficiency of aptamer attachment and the effectiveness of the washing steps, filtrates from washing after aptamer reaction were analyzed using a Nanodrop One C Microvolume UV-Vis Spectrophotometer (Thermo Scientific, USA). The concentration of the unbound aptamer in these filtrates was measured and compared to the initial aptamer stock solution.

The binding efficiency of the aptamer was determined by subtracting the cumulative aptamer in the collected filtrates from the initially added aptamer. Additionally, the final washing filtrate (fourth eluate) was analyzed to ensure the removal of any residual unbound aptamer.

### 2.9 Characterization of Aptamer Conjugated Nanoparticles

The size and surface charge of aptamer-conjugated NPs were measured using Malvern Zetasizer ZSP, as previously described. To further confirm aptamer attachment, gel electrophoresis (4% TBE) was performed. Free aptamer, conjugated nanoparticles, filtrates of purification, and non-modified nanoparticles were loaded onto the gel to visualize any shift in mobility, indicative of successful conjugation.

### 2.10 Evaluation of antibacterial efficacy *in vitro*

#### Bacteria preparation

*S. aureus* control strain ATCC 12600 was commercially obtained (KWIK STIK™, Microbiologics, USA), revived according to the manufacturer’s instructions, and stored as glycerol stocks at -80°C. The frozen bacterial cultures were grown for 16-18 hours at 35°C on Tryptic Soy Agar (TSA) plates and were further cultured in Tryptic Soy Broth (TSB) overnight for optimal growth. Before experiments, the bacterial cultures in TSB were spectrophotometrically assessed at 600 nm, and the colony-forming units/mL (CFU/mL) were enumerated using a standard growth curve and real-time microbial viability assay (BacTiter-Glo™, Promega, USA). The bacterial cultures were diluted to the required CFU/mL for subsequent antibacterial activity assays.

#### Minimum Inhibitory and Bactericidal Concentration (MIC and MBC) Assays

The antibacterial activity of Apt-P(V) NPs, P(V) NPs and free vancomycin was determined against *S. aureus* using a modified microbroth dilution method (CLSI protocol M07-A10). Briefly, ∼10^4^ CFU/mL of bacterial suspension prepared in cation-adjusted Mueller-Hinton Broth (Ca-MHB) was exposed to serial dilutions of prepared particles and free vancomycin formulations (stock concentration: 1-2 mg/mL and 0.1 mg/mL, respectively) in a sterile 96-well plate. The well plates were incubated at 35°C for 18-24 hours, and the turbidity was assessed both visually and spectrophotometrically (600 nm). The MIC was determined as the minimum concentration of particle/drug formulation that exhibited no visible turbidity. A 5µL drop from the clear wells was plated and incubated overnight on tryptic soy agar (TSA), and the MBC was determined as the minimum concentration that showed no viable colony growth. All experiments were performed in triplicate.

#### Minimum Biofilm Eradication Concentration (MBEC) Assay

Stainless steel (SS) plates (10 x 3 x 1mm) in sterile 24-well plates were exposed to ∼10^5^ CFU/mL of *S. aureus*, prepared in 1mL MHB broth for 6 hours at 35°C, to facilitate adhesion and biofilm formation on the surface. The spent media was discarded, and the SS plates were rinsed three times with 1x PBS, shaking for 2 minutes, to remove non-adherent bacteria. The SS plates were then treated with predetermined particle and drug formulations for 24 hours at 37°C. The coupons were subjected to a PBS wash cycle (2 mins, 3 times) and were sonicated for 40 minutes in 1mL PBS to retrieve the adherent bacteria. The bacterial viability in the sonicate fluid was determined in real-time by performing the BacTiter-Glo assay and capturing ATP-associated luminescence units using a microplate reader (Synergy H1, BioTek Instruments, USA). The sonicate fluid was also plated and incubated for 16-18 hours on TSA, and the viable bacteria were determined by the plate count method. MBEC was defined as the concentration of the formulation that resulted in a >3-log reduction in bacterial viability.

### 2.11 Aptamer Targeting Specificity and Biofilm Retention

#### Fluorescence Imaging for Aptamer Targeting Specificity

Sterile glass coverslips were placed in 6-well plates and preconditioned with 1 mL MHB media for 30 minutes. The media was aspirated and ∼10^5^ CFU/mL of *S. aureus*, prepared in 1mL MHB, were added to the wells containing the coverslips. The well plate was incubated for 1 hour at 35°C to facilitate bacterial adhesion. The spent media was replaced with fresh media, and the adhered bacteria were further incubated for 6 hours at 35°C to form biofilm on coverslips. The biofilms were then exposed to 200 µL of Cy5-tagged SA formulation for 1 hour at 35°C. The coverslips were washed 3 times with 1 mL PBS to remove any unbound particles, and the bacteria were stained with DAPI solution (1:1000 in DI water) for 15 minutes in the dark. The excess stain was washed off with 1 mL DI water (2x, 2 mins each). The coverslips were mounted using BacLight mounting oil (Thermo Fisher Scientific, USA), and fluorescent imaging was performed under the DAPI and Cy5 filters using a Nikon Eclipse TiE Microscope (100× magnification). Cy5-tagged versions of free SA aptamers, free scrambled aptamers, and Scr Apt-NPs served as controls. To assess non-specific aptamer attachment to the substrate, parallel coverslips without biofilm were incubated with the same Cy5-tagged aptamers and formulations and imaged under identical conditions.

#### Fluorescence Imaging for Nanoparticle Biofilm Retention

Rhodamine B (RhB) loaded PLGA NPs (P(RhB)) were conjugated with SA aptamers using EDC/NHS chemistry, as previously described in section 2.7. *S. aureus* biofilms were grown for 6 hours on sterile glass coverslips in 6-well plates as described above. After biofilm formation, 200□µL of P(RhB), with or without aptamer functionalization, were applied to each coverslip and incubated at 35□°C in the dark for 1 hour. Samples were washed three times with 1 mL of PBS (2 minutes each), stained with DAPI (1:1000) for 15 minutes, and then rinsed twice with PBS (1 mL, 2 minutes each). Coverslips were imaged using a Nikon Eclipse Ti-E fluorescence microscope (Nikon Instruments Inc., USA) equipped with a 60× objective and DAPI/TxRed filters (scale bar = 10□µm).

For quantification of biofilm retention, coverslips were incubated with NPs for 1 hour under the same conditions. After incubation, samples were washed to remove unbound particles and transferred to DMSO to dissolve retained nanoparticles. RhB fluorescence was measured using a Synergy H1 plate reader (BioTek Instruments, USA) at 540 nm excitation and 625 nm emission. Experiments were performed in triplicate, and fluorescence intensity was compared across treatment groups.

### 2.12 Retention and Biodistribution Study in the Rat Knee Joint

To evaluate intra-articular retention and systemic biodistribution, fluorescent SA Apt-NPs and fluorescent Scr Apt-NPs were prepared as described in section 2.7 using 3′-Cy5.5–tagged aptamers. These formulations were used to compare joint retention and biodistribution following intra-articular administration in infected and non-infected rat knee joints.

#### Animals and Study Timeline

All animal procedures were approved by the Massachusetts General Hospital Institutional Animal Care and Use Committee (protocol no. 2022N000013) and conducted in accordance with institutional and NIH guidelines for the care and use of laboratory animals. Male Sprague– Dawley rats (10–11 weeks old, 300–350 g; Charles River Laboratories, Wilmington, MA, USA) were housed under standard conditions with free access to chow and water. Animals were randomly assigned to infected and non-infected groups receiving either fluorescent SA Apt-NPs or fluorescent Scr Apt-NPs. Each group included n = 3 animals, unless otherwise indicated. Each rat received a single intra-articular injection of 50 µL of a 20 mg/mL NP suspension into the knee joint. For infected groups, nanoparticle injections were performed on postoperative day (POD) 7, following joint surgery and bacterial inoculation on POD 0.

#### Surgical Procedure and Infection Model

Rats were anesthetized with 1–3% isoflurane delivered in 1 L/min oxygen. Preoperative analgesia was provided using buprenorphine (0.05 mg/kg, intraperitoneally) administered 30 min before surgery. A 3–4 cm craniolateral skin incision was made over the right femur extending from the supracondylar region to the proximal tibia. The femoral condyle was exposed, and the overlying musculature was carefully dissected from the bone. A pilot hole (1 mm) was first drilled into the lateral femoral condyle, followed by enlargement to a 3 mm-diameter, 3 mm-deep cavity. A 3 × 3 mm cylindrical polyethylene (PE) plug (ethylene-oxide– sterilized, direct-compression-molded GUR ultrahigh-molecular-weight polyethylene without additives) was implanted transchondylarly into the femur. For infection induction, *S. aureus* (ATCC 12600, Manassas, VA, USA) was used [33]. An intercondylar canal was drilled in the proximal tibia, and 10 µL of bacterial suspension containing 2 × 10□CFU was injected into the canal, followed by implantation of a titanium maxillofacial screw (1.3 mm diameter × 8 mm length). Deep dermal closure was achieved using 5-0 Vicryl sutures, after which 40 µL of S. aureus suspension (8 × 10□CFU) was injected intraarticularly. The skin was closed using 5-0 Vicryl in a subcuticular fashion. Postoperative analgesia consisted of buprenorphine administered every 4–6 h for 72 h. Animals were euthanized at designated time points using pentobarbital (100 mg/kg, intraperitoneally) [19, 34–37].

#### IVIS Spectrum Imaging and Cy5.5 Quantification Analysis

Fluorescence retention in the joint was monitored in vivo using an IVIS Spectrum imaging system (PerkinElmer, USA) to track fluorescent SA Apt-NPs and Scr Apt-NPs carrying 3′-Cy5.5–tagged aptamers in infected and non-infected models. Imaging was performed immediately after injection and at 3, 6, 24, 48, and 72 hours, and weekly thereafter until the designated endpoint for each cohort. Fluorescence was quantified by drawing a constant-size region of interest (ROI) over the injected knee joint, with background subtraction using a non-fluorescent reference region. ROI size and placement were kept constant across all animals and timepoints. The signal from the injected knee was normalized to the contralateral knee for each animal and expressed as fold-change relative to the first post-injection time point. At the designated endpoint (POD35 or POD42, depending on the cohort), animals were euthanized, and major organs (liver, spleen, kidney, lung, heart, and brain) were harvested and imaged ex vivo under the same fluorescence settings (Cy5.5 excitation: ∼675 nm; emission: ∼720 nm).

#### Toe Spread

Gait parameters were quantified using ventral plane videography with the DigiGait™ Imaging System (Mouse Specifics, Inc., Framingham, MA, USA). Preoperative baseline recordings and postoperative assessments were acquired at a frame rate of 171 frames per second under standardized treadmill conditions. Toe spread was quantified from ventral plane gait recordings using ImageJ software (National Institutes of Health, Bethesda, MD, USA). Measurements were obtained by calculating the linear distance between digits 1-5 and 2-4, as visualized in representative frames. To account for inter-animal variability, values from the operated limb were normalized to those from the contralateral, non-operated control limb, and expressed as surgical-to-control ratios [38].

#### Static Weight-Bearing

Hindlimb weight distribution was assessed using a custom-built total internal reflection (TIR) imaging platform designed to capture plantar pressure profiles [37, 39]. Each rat was recorded continuously for 5 minutes under standardized conditions. From these recordings, eight distinct and stable hindlimb stance events per animal were extracted for analysis. Weight-bearing distribution ratios between the operated and contralateral control limbs were quantified using custom MATLAB scripts (MathWorks, Natick, MA, USA).

#### Ex Vivo Fluorescence Biodistribution Analysis

At the endpoint, animals were euthanized, and major organs and knee tissues were harvested for ex vivo fluorescence biodistribution analysis. Tissues were rinsed in PBS, weighed, and homogenized in lysis buffer consisting of PBS with Triton X-100 using a mechanical homogenizer. Homogenates were incubated on ice for 15–30 minutes and centrifuged at 12,000 × g for 15 minutes at 4°C to remove tissue debris. The supernatant containing extracted fluorescent signal was collected for analysis. A standard calibration curve was prepared by serially diluting Cy5.5 dye in the same lysis buffer, and 100 µL of each standard and sample were loaded into a black 96-well fluorescence plate in triplicate. Fluorescence was measured using a Synergy H1 plate reader (BioTek Instruments, USA) at excitation 675 nm and emission 694 nm. Cy5.5 concentrations were interpolated from the standard curve and normalized to tissue weight. Values at or below the background-corrected detection level were considered not detectable.

#### Histology Analysis

Spleen, liver, heart, lungs, brain, tibia, and kidneys were harvested and fixed in 4% neutral-buffered formalin for 24-48 hours. Tibial specimens subsequently underwent decalcification in 14% Versenate EDTA solution (American MasterTech Scientific, Lodi, CA, USA) at room temperature for 28 days with regular solution changes. Following fixation (and decalcification for bone), all tissues were processed through graded alcohols, cleared, and embedded in paraffin.

Coronal sections (5 µm thickness) spanning the fracture site were obtained from tibial samples, while corresponding sections were prepared from soft tissues. Sections were stained with hematoxylin and eosin (H&E) and digitized using a NanoZoomer Digital Pathology System (Meyer Instruments, Houston, TX, USA) at high resolution. Histological evaluation was performed qualitatively to assess cellular morphology, tissue organization, and distribution; no quantitative histomorphometric analyses were conducted.

### 2.13 Statistical Analysis

Data are presented as mean ± SD. Nanoparticle characterization, loading, release, and *in vitro* antibacterial/biofilm assays were analyzed using one-way or two-way ANOVA with Tukey’s post hoc test, based on experimental design, with at least n ≥ 3 independent experiments per condition. Toe spread and weight-bearing outcomes were compared between groups using a two-tailed Student’s t-test. (p < 0.05 was considered statistically significant.)

## 3 Results

### 3.1 Characterization of Vancomycin-loaded PLGA NPs

P(V) NPs were synthesized via a double emulsion flow-focusing microfluidic approach and exhibited a mean hydrodynamic diameter of 288.3 ± 22.2 nm and a zeta potential of −23.5 mV, as measured by dynamic light scattering (DLS) (Fig. 1a). Scanning electron microscopy (SEM) confirmed the spherical morphology of the lyophilized particles (Fig. 1b). The LC% of vancomycin was 7.9□±□2.2%, as determined from UV-Vis analysis of lyophilized particles. This direct quantification of washed, lyophilized particles ensured that only stably entrapped vancomycin contributed to the LC value, avoiding overestimation from loosely adsorbed or unencapsulated drug.

**Fig. 1.**
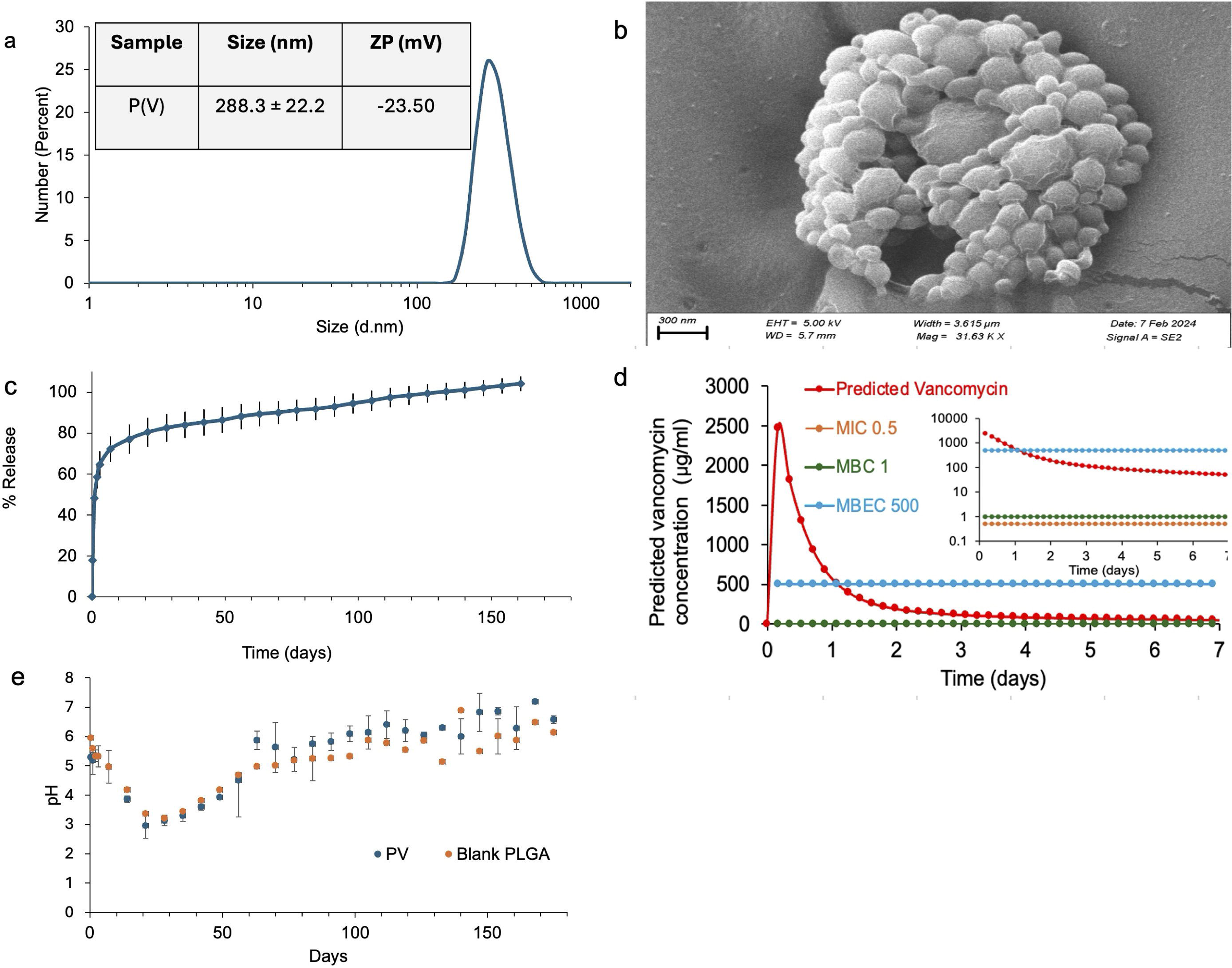
Characterization and release behavior of vancomycin-loaded PLGA nanoparticles. a) Representative DLS size distribution and zeta potential. b) SEM image showing nanoparticle morphology. c) Cumulative in vitro vancomycin release over 6 months. d) Predicted intra-articular vancomycin concentration over the first week after simulated administration of 160 mg P(V) NPs, modeled using first-order elimination with a 6 h half-life and 2 mL joint volume, shown relative to MIC, MBC, and the MBEC benchmark. The inset shows the same profile on a logarithmic y-axis to visualize lower concentration thresholds. e) pH changes in release eluates from vancomycin-loaded and blank PLGA nanoparticles over time.

### 3.2 In-vitro release profile

The in-vitro release profile of vancomycin from P(V) NPs showed an early high-release phase followed by an extended terminal release phase (Fig. 1c). Approximately 60% of the loaded vancomycin was released within the first 7 days, corresponding to the highest release rate during the early phase. Water-matched low-range UV–Vis validation gave an LOD of approximately 0.85 µg/mL and an LOQ of approximately 2.6 µg/mL. Based on interval release concentrations, vancomycin release remained above LOQ through approximately Day 21 and remained largely above LOD through approximately Day 161. The full release profile was therefore monitored through Day 180, with late time points interpreted in relation to the assay detection and quantification limits.

### 3.3 Pharmacokinetic (PK) modeling

In-vitro release model fitting to the cumulative data gave R² = 0.963 (SSE = 24.2) for the Korsmeyer–Peppas model, indicating that this model provided the best description of the release kinetics. Pharmacokinetic (PK) modeling based on the measured cumulative-release profile, scaled to 160 mg NPs in a one-compartment model with first-order elimination (t½ = 6 h, V = 2 mL), predicted an early burst with a maximum intra-articular concentration (Cmax) of 2.72 mg/mL at 1.31 h (Tmax). Predicted intra-articular concentrations were 0.051 mg/mL (51.0 µg/mL) at 1 week, 0.0276 mg/mL (27.6 µg/mL) at 2 weeks, and 0.0143 mg/mL (14.3 µg/mL) at 30 days. Corresponding AUC values were 13.83, 35.57, 43.55, and 54.15 mg·h/mL at 0.25, 1, 2, and 7 days, respectively, with a total AUC of 88.91 mg·h/mL over the monitored 182-day period.

Regarding the activity of vancomycin against S. aureus (MIC = 0.5 µg/mL; MBC = 1 µg/mL; MBEC benchmark = 500 µg/mL), model-based estimates indicated that predicted intra-articular concentrations remained above the measured MIC and MBC throughout the monitored 182-day release period. The predicted concentration exceeded the MBEC benchmark for approximately 1.1 days (∼26 h), supporting an early high-exposure phase. Because terminal interval release approached the assay quantification range, late-duration estimates were interpreted cautiously and used to contextualize the extended release profile rather than to define precise therapeutic exposure at late time points.

### 3.4 PLGA Degradation as Indicated by pH of Release Medium

The pH of the release medium was monitored over 6 months for both vancomycin-loaded and blank NPs as an indirect indicator of PLGA degradation-associated acidification. Both formulations showed an initial decrease in pH during the first 10–15 days (Fig. 1e). The vancomycin-loaded nanoparticles exhibited a slightly lower starting pH than blank nanoparticles, likely reflecting the contribution of the encapsulated vancomycin formulation and/or residual acidic components. After this early decrease, the pH of both formulations gradually increased toward the initial range by approximately Day 60–90 and remained relatively stable thereafter. This pattern suggests that the major acidification phase occurred during the early-to-intermediate release period, while later release measurements represented an extended terminal phase rather than evidence of a separate prolonged degradation-controlled release phase.

### 3.5 Aptamer Conjugation and Attachment Efficiency

Nanoparticles conjugated with biofilm-specific aptamers (SA Apt-P(V)) via EDC/NHS chemistry exhibited a hydrodynamic diameter of 307.9 ± 10.84 nm and a zeta potential of –47.49 mV (Fig. 2). UV–Vis absorbance measurements of washing eluates confirmed the absence of detectable free aptamer in the final wash (Data not shown). The absence of free aptamer and the retention of aptamer-conjugated nanoparticles during electrophoresis further supported successful conjugation and formulation stability (Supplementary Fig. 2).

**Fig. 2.**
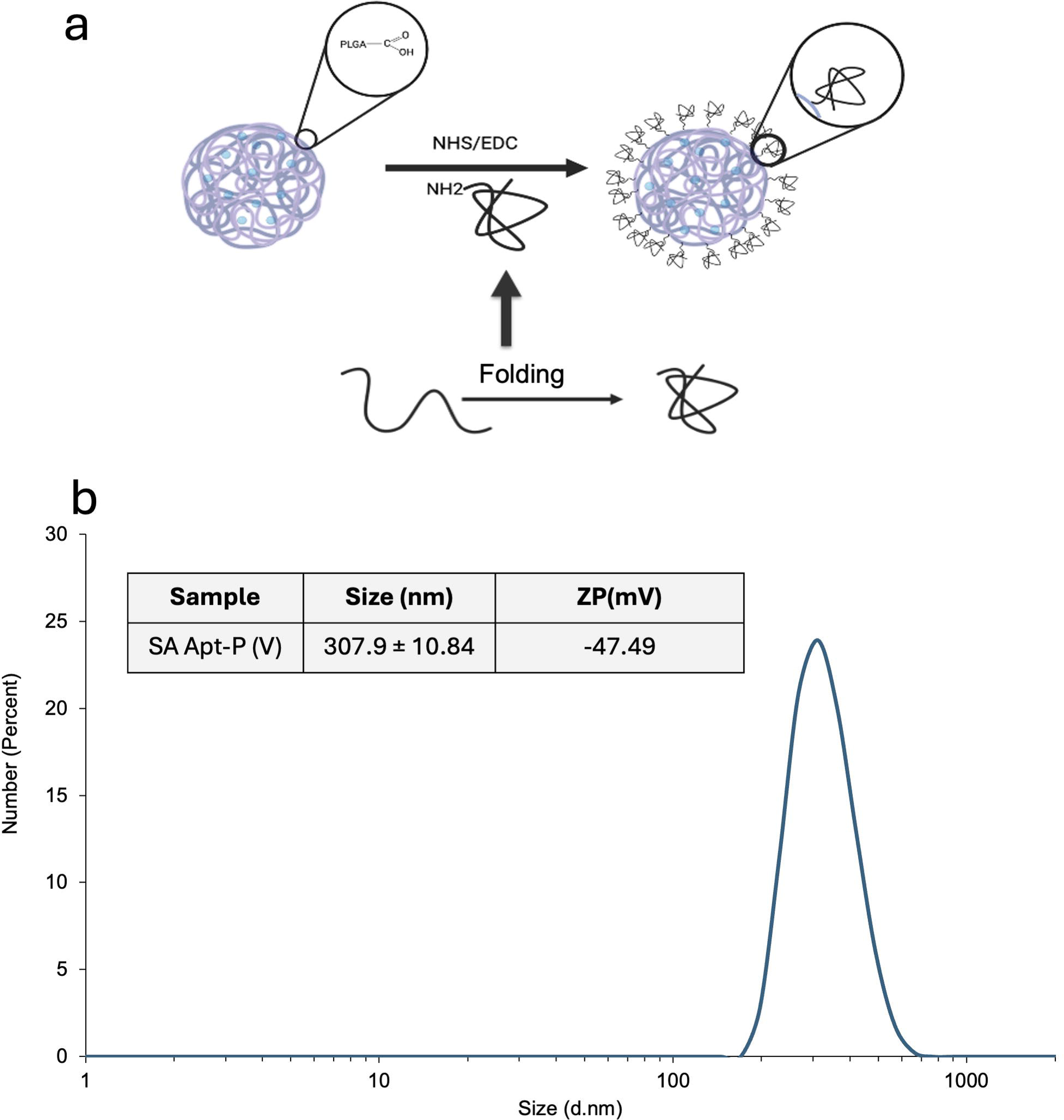
SA Aptamer conjugation to PLGA nanoparticles. a) Schematic of SA-specific aptamer folding and EDC/NHS-mediated conjugation to PLGA nanoparticles. Aptamers were folded by heating at 95°C for 5 min and cooling on ice for 15 min before conjugation. b) Representative DLS size distribution and zeta potential of aptamer-conjugated PLGA nanoparticles, showing a hydrodynamic diameter of 307.9 ± 10.84 nm and zeta potential of −47.49 mV. Gel electrophoresis confirmation is shown in Supplementary Fig. 2.

### 3.6 Antibacterial efficacy *in vitro*

The MIC and MBC for free vancomycin were 0.75±0.3 and 1.5±0.5 µg/mL, respectively. The MIC of particle formulations P(V) and SA Apt-P(V) were 20.8±7.3 and 39±12.8 µg/mL, and MBC were 142±52 and 175±43.3 µg/mL, respectively, with no significant differences in their activity (Table 2).

**Table 2:** Minimum inhibitory and bactericidal concentrations of prepared vancomycin-loaded particles and free vancomycin.

| loaded particles and free vancomycin |  |  |
| --- | --- | --- |
|  | MIC (µg/mL) | MBC (µg/mL) |
| PLGA(V) | 20.8±7.3 | 142±52 |
| SA Apt-PLGA(V) | 39±12.8 | 175±43.3 |
| Free Vancomycin | 0.75±0.3 | 1.5±0.5 |

The antibiofilm activity of vancomycin nanoparticles was evaluated by determining MBEC (Fig. 3). The P(V) NPs demonstrated biofilm eradication at all tested concentrations (1× to 10× MIC). The SA Apt-P(V) particles showed biofilm eradication consistent with the P(V) particles in real time (BacTiter-Glo™ assay), with some recovery of viable bacteria observed in the plate count method for all concentrations except 10×MIC. Control P(V) particles (cP(V)) underwent the same post-processing workflow as SA Apt-P(V) but without aptamer conjugation. cP(V) showed lower antibiofilm activity than P(V), whereas SA Apt-P(V) showed greater bacterial reduction than cP(V) despite undergoing the same post-processing workflow and containing a comparable vancomycin payload. Free vancomycin (FV) produced variable reductions in bacterial viability across concentrations (1×–100× MIC) in the BacTiter Glo™ assay; however, viable bacteria were recovered by plate counting at all tested concentrations, indicating limited antibiofilm activity. The bacterial viability from biofilms without antibiotic exposure was determined in the range of 10^4^ – 10^5^ CFU/mL.

**Fig. 3.**
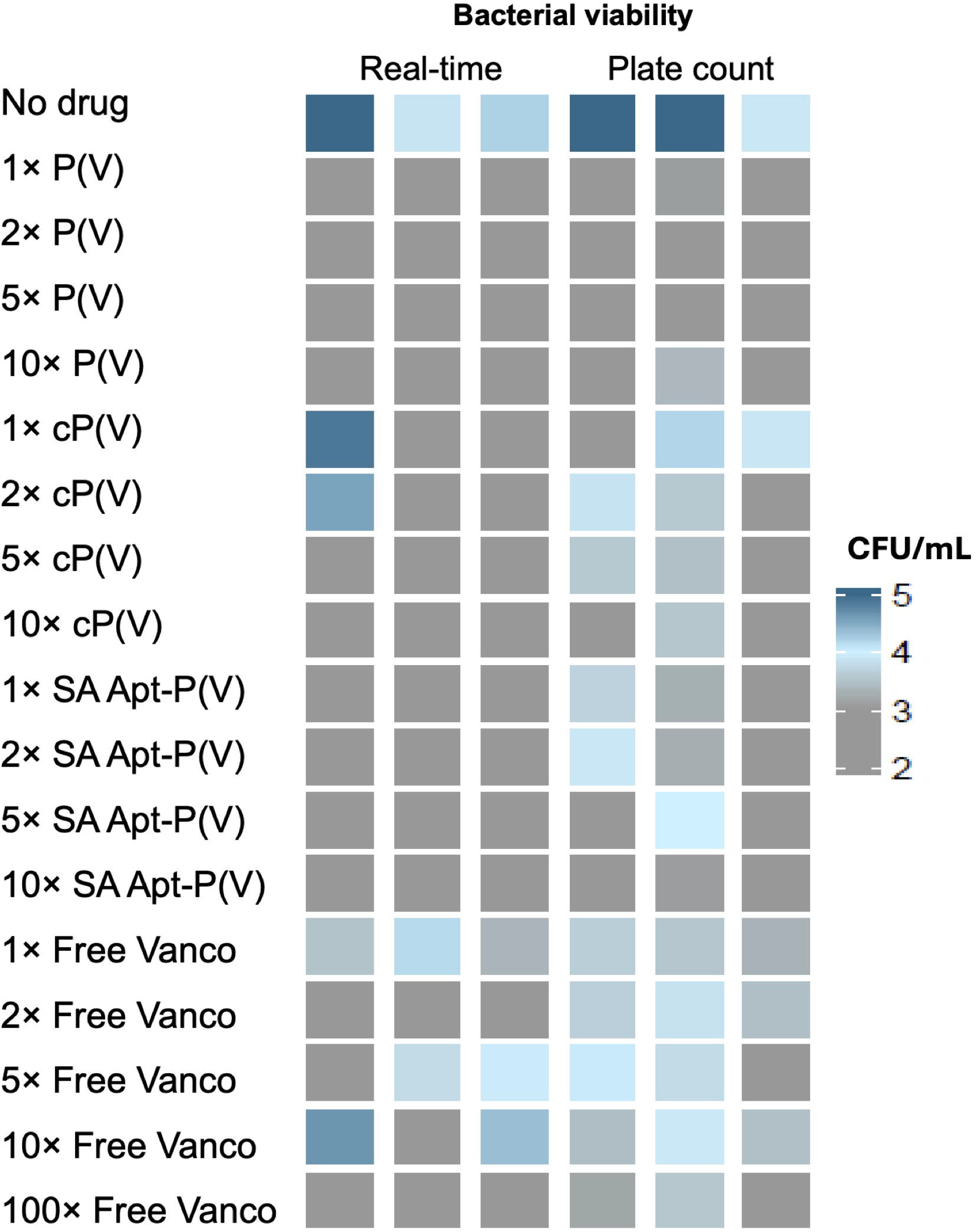
Vancomycin-loaded nanoparticle formulations show antibiofilm activity against S. aureus biofilms. Heatmap showing recovered adherent bacterial viability after treatment of S. aureus biofilms grown on stainless steel coupons with P(V), cP(V), SA Apt-P(V), or free vancomycin. cP(V) refers to the processed control formulation that underwent the same post-processing workflow as SA Apt-P(V) but without aptamer conjugation. Columns represent real-time ATP-based microbial viability assay and plate count, performed in triplicate. The color scale represents log_₁₀_ CFU/mL. Biofilms without drug exposure served as the no-drug control.

### 3.7 Fluorescence Imaging for Aptamer Targeting Specificity

#### Visualization of biofilm-targeting properties of S. aureus-specific Apt-NPs

Cy5 fluorescence from fluorescent SA Apt-NPs was observed to localize on and around DAPI-stained bacterial aggregates embedded within the biofilm matrix (Fig. 4). Cy5-tagged free SA aptamers served as positive control and showed strong colocalization with the bacteria. In contrast, no significant Cy5 signal was observed in biofilms treated with fluorescent Scr Apt-NPs or Cy5-tagged free scrambled aptamers, which served as negative controls.

**Fig. 4.**
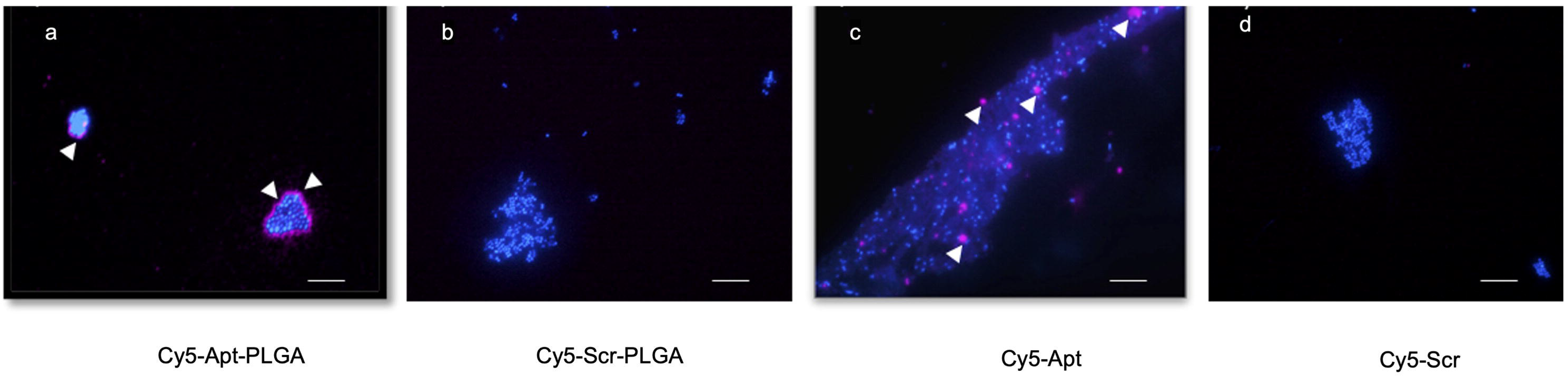
Fluorescence microscopy of aptamer-mediated targeting of S. aureus biofilms. a) Cy5-SA Apt-NPs incubated with DAPI-stained nascent 6 h S. aureus biofilms grown on glass coverslips. b) Cy5-Scr Apt-NPs. c) Free Cy5-labeled SA aptamer, used as a positive control. d) Free Cy5-labeled scrambled aptamer, used as a negative control. Blue indicates DAPI-stained bacteria/biofilm, and magenta indicates Cy5 signal. White arrows indicate aptamer-associated binding on and around S. aureus biofilms. Scale bar = 10 µm.

#### Fluorescence Imaging for Nanoparticle Biofilm Retention

To evaluate biofilm retention, non-targeted P(RhB) NPs and targeted SA Apt-P(RhB) NPs were applied to biofilms and imaged after DAPI staining. Untreated biofilms showed no detectable fluorescence signal (Fig. 5a). Biofilms treated with non-targeted P(RhB) NPs exhibited minimal RhB signal (Fig. 5b). In contrast, biofilms exposed to targeted SA Apt-P(RhB) NPs showed markedly higher RhB fluorescence, indicating enhanced retention within the biofilm matrix (Fig. 5c). Quantitative analysis using DMSO extraction and fluorescence measurement at 540 nm excitation and 625 nm emission confirmed significantly greater RhB retention for SA Apt-P(RhB) NPs (276.1 ± 20.6 a.u.) compared with non-targeted P(RhB) NPs (98.9 ± 3.7 a.u.) (p < 0.01). Minimal fluorescence was detected in MgCl_₂_-treated biofilms (0.00 ± 0.14 a.u.) and untreated control biofilms (5.35 ± 2.40 a.u.), confirming the observed signal resulted from nanoparticle retention rather than background fluorescence.

**Fig. 5.**
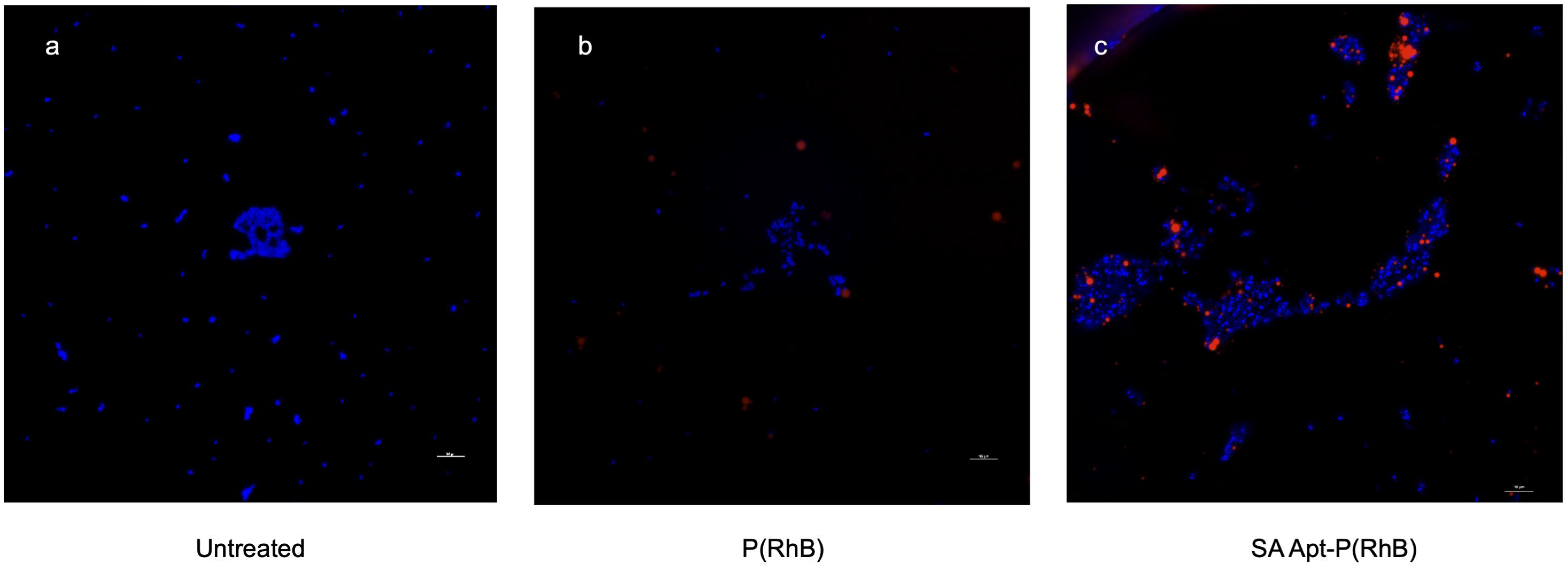
Aptamer-mediated retention of RhB-loaded nanoparticles in S. aureus biofilms. Fluorescence microscopy images of DAPI-stained biofilms: a) untreated biofilm control, b) biofilm treated with non-targeted P(RhB) nanoparticles, and c) biofilm treated with SA Apt-P(RhB) nanoparticles. Blue indicates DAPI-stained bacteria/biofilm, and red indicates RhB fluorescence. Scale bar = 10 µm.

### 3.8 Retention and Biodistribution in the Rat Knee Joint

Quantitative IVIS analysis of knee joint fluorescence (Fig. 6) showed prolonged retention of fluorescent aptamer-conjugated nanoparticles following intra-articular administration. Fluorescent SA Apt-NPs exhibited higher retention in infected joints compared with non-infected joints over the early post-injection period (Fig. 6a). In the infected model, fluorescent SA Apt-NPs showed higher mean fluorescence than fluorescent Scr Apt-NPs, although inter-group differences decreased over time as the signal declined (Fig. 6b). Despite the shorter 28-day post-injection follow-up for the infected groups compared with 42 days for the non-infected group, cumulative AUC was higher in both infected groups, with the highest AUC observed for SA Apt-NP infection (11.83), followed by Scr Apt-NP infection (6.69) and SA Apt-NP non-infection (0.74) (Table 3).

**Fig. 6.**
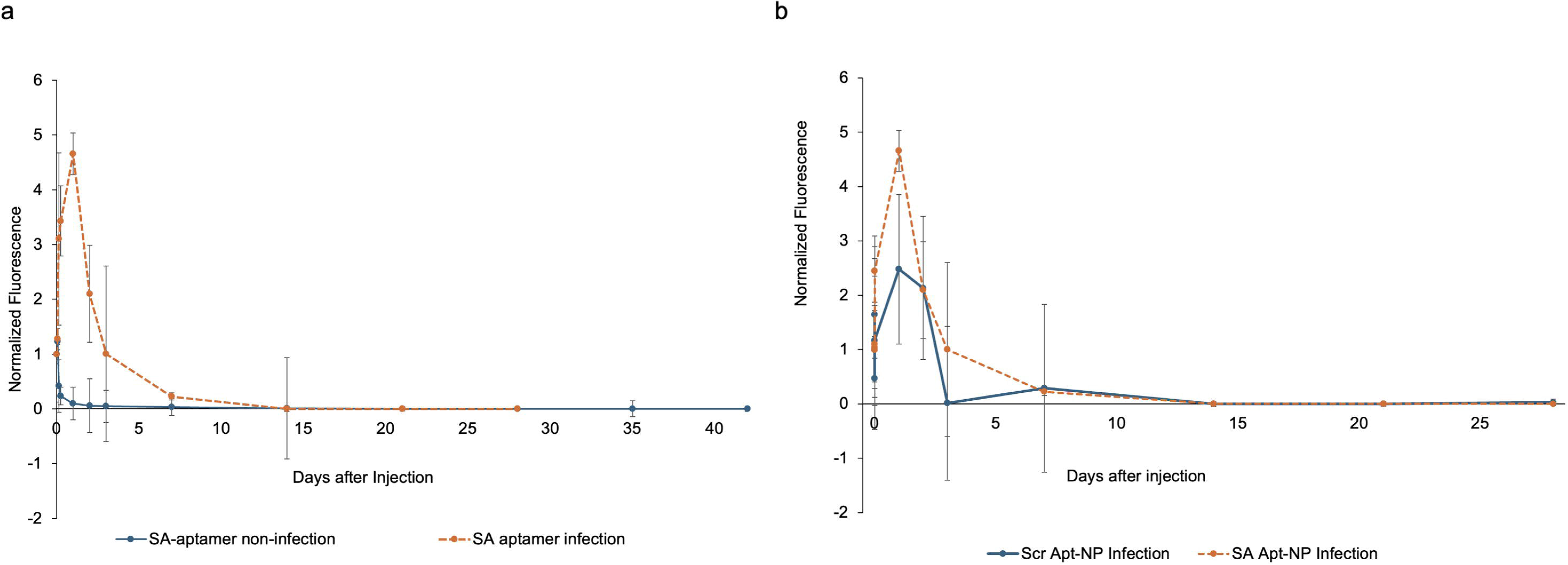
In vivo fluorescence retention of Cy5.5-labeled aptamer-conjugated nanoparticles after intra-articular administration. a) Retention of SA Apt-NPs in infected versus non-infected joints over time. b) Retention of SA Apt-NPs compared with Scr Apt-NPs in the infection model. Fluorescence was quantified using a constant-size region of interest over the injected knee joint, background-subtracted, normalized to the contralateral knee, and expressed as fold-change relative to the first post-injection time point for each animal. Data are shown as mean ± SD.

**Table 3.** Area-under-the-curve (AUC) analysis of fluorescence retention efficiency following intra-articular administration of Cy5.5-labeled nanoparticles.

| Rat Group | SA Apt-NP<br>Non-Infection | Scr Apt-NP<br>Infection | SA Apt-NP<br>Infection |
| --- | --- | --- | --- |
| Days after injection | 42 | 28 | 28 |
| Efficiency AUC<sub>0-days after injection</sub> | 0.74 | 6.69 | 11.83 |

*AUC values were calculated from longitudinal IVIS measurements of retention efficiency over the available post-injection follow-up period for each group (42 days for SA Apt-NP non-infection and 28 days for both infected groups). Higher AUC values indicate greater cumulative nanoparticle retention over the corresponding observation period.*

#### Toe Spread Ratio

To assess functional outcomes in the infected model, the SA Apt-NP and Scr Apt-NP groups were combined and compared with the previously published infection-control group and infection database [19, 34]. Toe spread measurements in the combined aptamer-treated group decreased immediately after surgery and gradually returned to baseline by postoperative day (POD) 14, consistent with previous data [19]. For the 1-5 toe spread ratio, the combined aptamer-treated group demonstrated a significant reduction on POD 1, 3, 7, 21, 28, and 35 compared with preoperative values (p < 0.05). For the 2-4 toe spread ratio, the combined aptamer-treated group showed a significant reduction on POD 1 and 3, relative to preoperative values (p < 0.05) (Fig. 7).

**Fig. 7.**
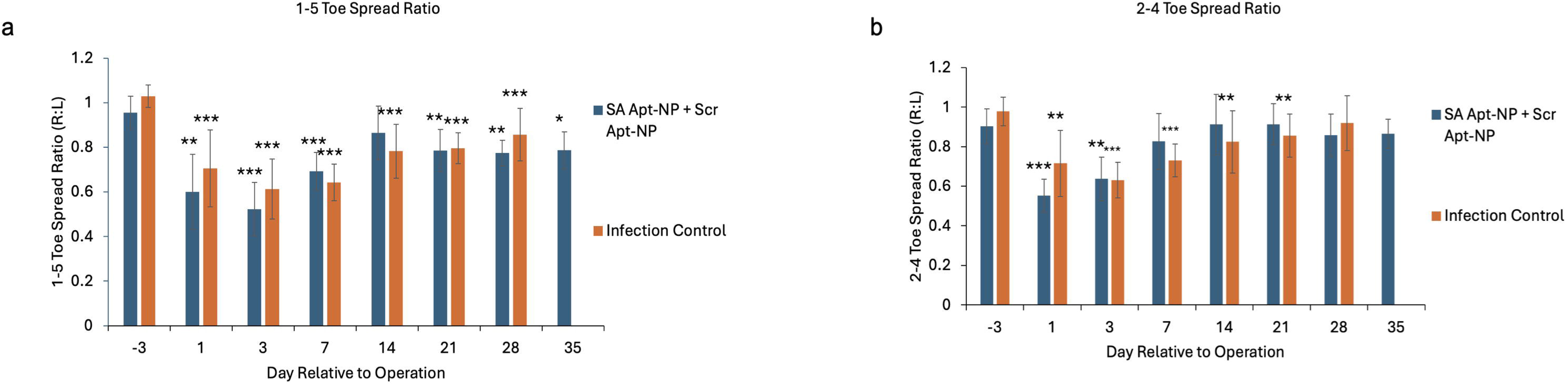
Toe-spread analysis following intra-articular administration of aptamer-conjugated nanoparticles. Functional outcome assessment showing hind paw right:left surgical:non-surgical toe-spread ratio. a) 1–5 toe-spread ratio. b) 2–4 toe-spread ratio. Data are shown as mean ± SD. Asterisks indicate significant differences compared with preoperative baseline or between groups. *p < 0.05, **p < 0.01, ***p < 0.001, #p < 0.05.

#### Weight-Bearing Ratio

Weight-bearing assessments demonstrated that both the SA Apt-NP group and the previously published infection-control group exhibited reductions immediately after surgery, which did not return to baseline during the observation period. The SA Apt-NP group demonstrated significant reductions on POD 14, 21, and 35 (p < 0.05). A significant difference between the SA Apt-NP group and the previously published infection-control group was observed at POD 7 (p < 0.05) (Fig. 8).

**Fig. 8.**
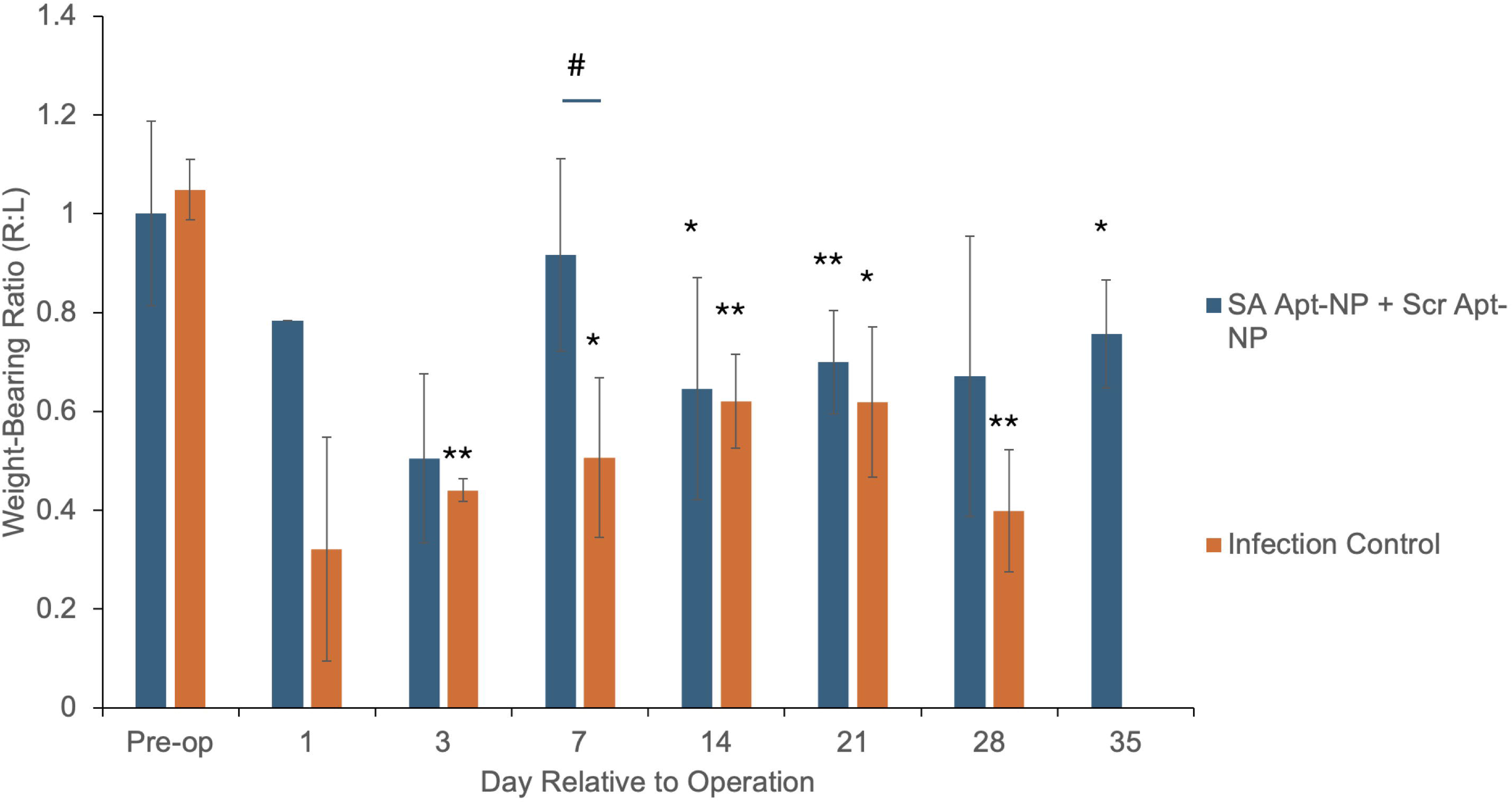
Static weight-bearing analysis following intra-articular administration of aptamer-conjugated nanoparticles. Weight-bearing right:left surgical: non-surgical ratio for hind limbs over time. Data are shown as mean ± SD. Asterisks indicate significant differences compared with preoperative baseline or between groups. *p < 0.05, **p < 0.01, #p < 0.05.

#### Ex Vivo Fluorescence Biodistribution Analysis

Endpoint ex vivo imaging of harvested tissues showed no appreciable fluorescence signal in major off-target organs, including spleen, liver, heart, lung, brain, and kidney (Supplementary Fig. 3). Plate-reader quantification of extracted Cy5.5 from tissue homogenates was at or near background levels in these organs, supporting the absence of detectable long-term off-target tissue accumulation. Representative tissue images are shown in Supplementary Fig. 3 to document the biodistribution pattern at the endpoint.

#### Histology Analysis

Histological examination of major organs did not reveal evidence of nanoparticle administration– associated systemic pathological alterations across the experimental groups (Fig. 9).

**Figure 9.**
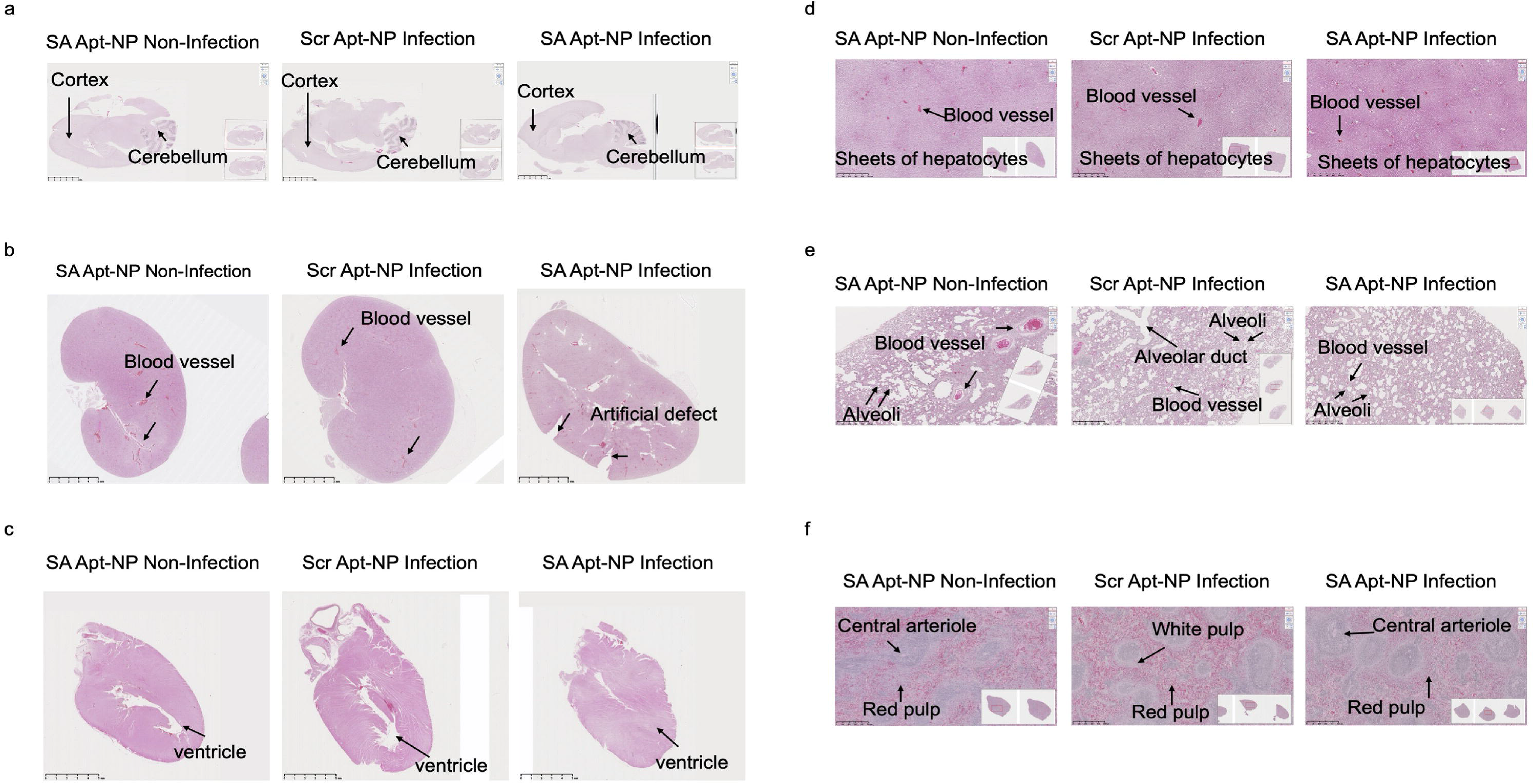
Histological evaluation of major organs following intra-articular nanoparticle administration. H&E-stained sections of a) brain, b) kidney, c) heart, d) liver, e) lung, and f) spleen following intra-articular nanoparticle administration. Scale bars: 5 mm in panels a–c and 500 µm in panels d–f. No overt inflammation or necrosis was observed in the examined organs.

In the brain, normal cytoarchitecture was preserved in both the cortex and cerebellum, with intact neuronal layering and no signs of neuronal degeneration, gliosis, hemorrhage, or inflammatory cell infiltration. The kidney exhibited well-defined glomeruli and renal tubules with preserved basement membranes and no evidence of tubular necrosis, interstitial inflammation, or vascular congestion.

Cardiac sections demonstrated normal myocardial fiber organization, intact ventricular architecture, and absence of inflammatory infiltrates, fibrosis, or cardiomyocyte degeneration.

The liver showed preserved hepatic lobular architecture with orderly sheets of hepatocytes and intact sinusoidal structures. No hepatocellular necrosis, steatosis, inflammatory infiltration, or sinusoidal dilation was observed.

Lung sections displayed normal alveolar architecture with thin alveolar septa, clear alveolar spaces, and intact pulmonary vasculature, without evidence of edema, hemorrhage, or inflammatory cell accumulation.

In the spleen, both red pulp and white pulp regions were well organized, with clearly identifiable central arterioles and no signs of architectural disruption, lymphoid depletion, or necrosis.

In the tibia, H&E histology confirmed infection-associated tissue changes in both Apt-NP– treated infected groups. Qualitatively, SA Apt-NP injection did not show overt worsening of inflammatory or destructive histopathology compared with scrambled Apt-NP injection. A previously published PJI-only histologic image was included as a historical reference only and was not part of the current study or formal comparison (Supplementary Fig. 4).

## 4. Discussion

PJI remains difficult to treat because biofilm-associated bacteria are poorly exposed to systemic antibiotics, while sustained high local antibiotic concentrations are challenging to achieve without toxicity. To address this gap, we developed vancomycin-loaded PLGA nanoparticles and conjugated them with SA-specific aptamers to improve biofilm localization and retention (Figs. 1–2).

Encapsulation of hydrophilic vancomycin into hydrophobic PLGA remains challenging because of weak drug–polymer interactions. In our unpublished manual W/O/W formulation experiments using comparable polymer–drug ratios and the same direct LC quantification workflow, loading capacities were typically below 1%, with occasional batches reaching 2–3% but with high variability, consistent with the difficulty reported in prior studies [17, 18, 20]. In contrast, the microfluidic formulation used here achieved higher vancomycin loading, which is important because therapeutic dose is determined by the final drug content in purified particles rather than encapsulation efficiency alone.

Chip-to-chip and lot-to-lot variability in loading nevertheless remained a limitation. This likely reflects the sensitivity of flow-focusing microfluidic synthesis to subtle differences in channel morphology, surface properties, and mixing behavior, which can alter nanoprecipitation dynamics for hydrophilic and bulky drugs such as vancomycin [40–43]. These findings emphasize that improving loading robustness and implementing chip qualification and process controls will be essential for translation. The nanoscale size of these particles provides an additional advantage for biofilm-associated infection, where smaller particles can more effectively interact with biofilm matrices than micron-scale depots [44, 45].

Release studies showed an early high-release phase followed by slower extended release, with interval release interpreted in relation to the water-matched analytical detection and quantification limits (Fig. 1c). To relate these findings to pharmacodynamic benchmarks, we modeled intra-articular vancomycin concentrations assuming administration of 160 mg total PLGA mass, corresponding to the carrier mass of the FDA-approved intra-articular PLGA formulation Zilretta® [46]. Assuming first-order clearance of vancomycin based on a half-life of 6 hours [47], model-based estimates suggested that predicted intra-articular concentrations remained above the measured MIC and MBC throughout the monitored 182-day release period, although late-duration estimates were interpreted cautiously because terminal interval release approached the assay quantification range. Concentrations exceeded the MBEC benchmark (500 µg/mL) for approximately 1.1 days (∼26 h) during the early high-exposure phase. The modeled 24-h AUC/MIC also exceeded the systemic therapeutic target commonly referenced for intravenous vancomycin therapy [48]. Although MBEC values are assay-dependent and do not represent formal intra-articular pharmacodynamic targets, these findings suggest that the formulation can provide an early antibiofilm exposure phase followed by prolonged maintenance above MIC. Based on prior work defining therapeutic goals for nascent implant-associated infections [36, 49], these results support the feasibility of sustained local vancomycin delivery using PLGA nanoparticles.

Monitoring of release eluate pH provided an indirect indicator of PLGA degradation-associated acidification (Fig. 1e). Both vancomycin-loaded and blank PLGA nanoparticles showed an early pH decrease during the first 10–15 days, consistent with PLGA hydrolysis and accumulation of acidic degradation products [15, 50].

The pH then gradually recovered toward the initial range by approximately Day 60–90, consistent with progressive neutralization/removal of acidic byproducts as the system evolved over time [51].

Therefore, the 180-day release profile was interpreted as an early high-release phase followed by extended terminal release, rather than as evidence of a distinct prolonged degradation-controlled phase. Late time points were interpreted in relation to the assay detection and quantification limits.

Aptamer conjugation was supported by changes in hydrodynamic size, zeta potential (Fig. 2), and electrophoresis patterns (Supplementary Fig. 2). Functionally, SA aptamer-conjugated nanoparticles showed stronger association with S. aureus biofilms than scrambled or non-targeted controls (Fig. 4). RhB-loaded targeted particles also showed greater retention within biofilms than non-targeted particles (Fig. 5), indicating that aptamer conjugation improved cargo localization. This targeting effect was consistent with the in-vitro antibacterial results, where aptamer-conjugated vancomycin-loaded particles improved antibiofilm performance compared with non-targeted controls at equivalent vancomycin concentrations (Fig. 3; Table 2).

In-vivo imaging was used to evaluate retention, biodistribution, and preliminary biocompatibility after intra-articular administration in the rat PJI model [34]. Fluorescent nanoparticles showed prolonged residence within the knee joint, while endpoint tissue analysis showed minimal signal in major organs, supporting limited long-term off-target tissue accumulation. In the infection model, SA Apt-NPs showed higher mean joint fluorescence than scrambled controls, although the difference between targeted and scrambled particles decreased over time as the signal declined. Endpoint histological analysis showed no evidence of treatment-associated tissue injury, supporting a favorable preliminary safety profile following intra-articular administration.

At the formulation level, batch-to-batch differences in microfluidic chip morphology introduce variability in drug loading; hydrophilic drug encapsulation also remains sensitive to small shifts in solvent conditions. In addition, we observed loss of drug during aptamer attachment and subsequent purification steps, which reduces the effective payload of the conjugated formulation relative to the control nanoparticles. Biofilm assays *in vitro*, while essential for screening, are limited by variability in biofilm density and composition and do not fully recapitulate host-conditioned matrices. In animal studies, increasing sample size may provide more statistical power to detect subtle differences in gait coordination and functional metrics. Biodistribution was assessed at the tissue level; more quantitative approaches (e.g., microdialysis or radiolabeling) could provide higher-resolution pharmacokinetic data within the joint.

Importantly, this study did not evaluate antibacterial efficacy *in vivo*. The present work was designed to establish formulation performance, biofilm targeting, biodistribution, and safety following intra-articular administration. Because aptamer conjugation reduced the final drug payload and introduced formulation variability, further optimization of payload retention and dose matching will be needed before evaluating therapeutic efficacy *in vivo*.

## 5. Conclusions

Taken together, this study demonstrates that SA aptamer-conjugated PLGA nanoparticles can combine sustained local vancomycin delivery with improved biofilm localization. The formulation achieved higher vancomycin loading than typical manual W/O/W preparations, provided an early high-exposure release phase followed by extended release, and showed improved biofilm interaction *in vitro* and antibiofilm activity after aptamer conjugation. In-vivo imaging supported localized joint retention with minimal long-term off-target tissue accumulation, and histology showed no overt systemic toxicity at the tested time point. Overall, this platform supports further development of targeted intra-articular nanoparticle delivery strategies for PJI, with future studies needed to test therapeutic efficacy *in vivo*.

## List of abbreviations

AUC: Area under the curve
Ca-MHB: Cation-adjusted Mueller-Hinton broth
CFU: Colony-forming units
C_max_: Maximum concentration
Cy5: Cyanine 5
Cy5.5: Cyanine 5.5
DAPI: 4′,6-diamidino-2-phenylindole
DAIR: Debridement, antibiotics, and implant retention
DLS: Dynamic light scattering
DMSO: Dimethyl sulfoxide
EDC: 1-ethyl-3-(3-dimethylaminopropyl)carbodiimide
FV: Free vancomycin
H&E: Hematoxylin and eosin
IACUC: Institutional Animal Care and Use Committee
IVIS: In vivo imaging system
LC: Loading capacity
LOD: Limit of detection
LOQ: Limit of quantification
MBC: Minimum bactericidal concentration
MBEC: Minimum biofilm eradication concentration
MHB: Mueller-Hinton broth
MIC: Minimum inhibitory concentration
MRSA: Methicillin-resistant Staphylococcus aureus
MSSA: Methicillin-sensitive Staphylococcus aureus
MWCO: Molecular weight cutoff
NHS: N-hydroxysuccinimide
NPs: Nanoparticles
PBS: Phosphate-buffered saline
PE: Polyethylene
PJI: Prosthetic joint infection
PK: Pharmacokinetic
PLGA: Poly(lactic-co-glycolic acid)
POD: Postoperative day
PVA: Polyvinyl alcohol
RCF: Relative centrifugal force
RhB: Rhodamine B
ROI: Region of interest
SA: Staphylococcus aureus
SD: Standard deviation
SEM: Scanning electron microscopy
SS: Stainless steel
TBE: Tris-borate-EDTA
TIR: Total internal reflection
Tmax: Time to maximum concentration
TSA: Tryptic soy agar
TSB: Tryptic soy broth
UV–Vis: Ultraviolet–visible spectroscopy
W/O/W: Water-in-oil-in-water

## Declarations

### Ethics approval and consent to participate

All animal procedures were approved by the Institutional Animal Care and Use Committee (IACUC; protocol no. 2022N000013) and conducted in accordance with institutional and NIH guidelines for the care and use of laboratory animals.

### Consent for publication

Not applicable.

### Availability of data and materials

All data generated or analyzed during this study are included in this article and its supplementary information files. Additional data are available from the corresponding author upon reasonable request.

### Competing interests

The authors declare no competing financial interest.

### Funding

This study was funded by the Muratoglu-Harris Endowment.

## Authors’ contributions

Conceptualization and study design: ZE, AS, YF, and EO. Nanoparticle formulation, drug loading, release studies, aptamer conjugation, nanoparticle characterization, and biofilm-targeting experiments: ZE. In vitro antibacterial and biofilm assays: AS, ZE, and KF. Animal procedures, functional outcome analysis, biodistribution assessment, and histological evaluation: YF, KF, PET, FBM, and YY. Pharmacokinetic modeling and microfluidic device support: MDA. Data analysis and interpretation: ZE, YF, AS, and EO. Manuscript drafting: ZE, YF, and AS. Manuscript review and editing: ZE and EO. Scientific supervision and resources: OM and EO. All authors read and approved the final manuscript.

## Supporting Information

Supplementary experimental details and figures describing the microfluidic nanoparticle preparation, aptamer-conjugation validation, ex vivo biodistribution, and historical reference histology.

## Acknowledgements

Part of this work was performed at the Center for Nanoscale Systems (CNS), Harvard University, a member of the National Nanotechnology Coordinated Infrastructure Network (NNCI), supported by NSF award ECCS-2025158. During the preparation of this work, the authors used ChatGPT (OpenAI) for limited language refinement and editorial assistance, and BioRender AI (BioRender) to assist with creation of the graphical abstract. After using these tools, the authors reviewed and edited the content as needed and take full responsibility for the content of the published article.

## Supplementary figure titles and legends

Supplementary Fig. 1. Design of the dual flow-focusing microfluidic device used for nanoparticle synthesis. a) Two-dimensional schematic of the microfluidic channel layout illustrating the dual flow-focusing geometry and key channel dimensions used to generate W/O/W emulsions. b) Three-dimensional rendering of the microfluidic device showing inlet and outlet ports and overall device architecture.

Supplementary Fig. 2. Gel electrophoresis confirmation of aptamer conjugation. Lane 1: DNA ladder; lane 2: SA Apt-NPs; lane 3: free SA aptamer, 1 µL; lane 4: first filtrate, 1 µL; lane 5: first filtrate, 10 µL; lane 6: second filtrate, 10 µL; lane 7: third filtrate, 10 µL; lane 8: unconjugated NPs. Arrow indicates the expected 88-nt aptamer band. The gel supports aptamer association with nanoparticles and removal of unbound aptamer during purification.

Supplementary Fig. 3. Representative ex vivo IVIS images of harvested tissues at endpoint. Representative ex vivo IVIS images of harvested tissues from SA Apt-NP non-infected, Scr Apt-NP infected, and SA Apt-NP infected groups. Images include major off-target organs and knee tissues. These representative images document the absence of appreciable endpoint fluorescence signal in major off-target organs.

Supplementary Fig. 4. Histologic evaluation of infected joints following Apt-NP injection with historical PJI reference histology. Representative H&E-stained histologic sections from infected joints treated with scrambled Apt-NP or Staphylococcus aureus–targeted Apt-NP are shown in cross-sectional and anteroposterior (AP)-oriented views. Both Apt-NP–treated infected groups demonstrated periarticular inflammatory tissue formation and capsule-like remodeling, supporting successful establishment of the infection model. Qualitative histologic assessment did not show overtly increased inflammatory tissue expansion, suppurative change, necrosis, or structural tissue destruction in the S. aureus–targeted Apt-NP group compared with the scrambled Apt-NP group. A representative PJI-only histologic section from a previously published rat PJI model [34] is included as a historical reference to provide visual context for infection-associated histopathology. Arrows indicate infection-associated periarticular inflammatory tissue/capsule formation. Scale bars= 2.5 mm; 5mm.

## Notes

### Competing Interest Statement

The authors have declared no competing interest.

