## Supplementary Figures for "Aptamer-Targeted PLGA Nanoparticles for Biofilm-Directed Intra-articular Antibiotic Delivery in Prosthetic Joint Infection"

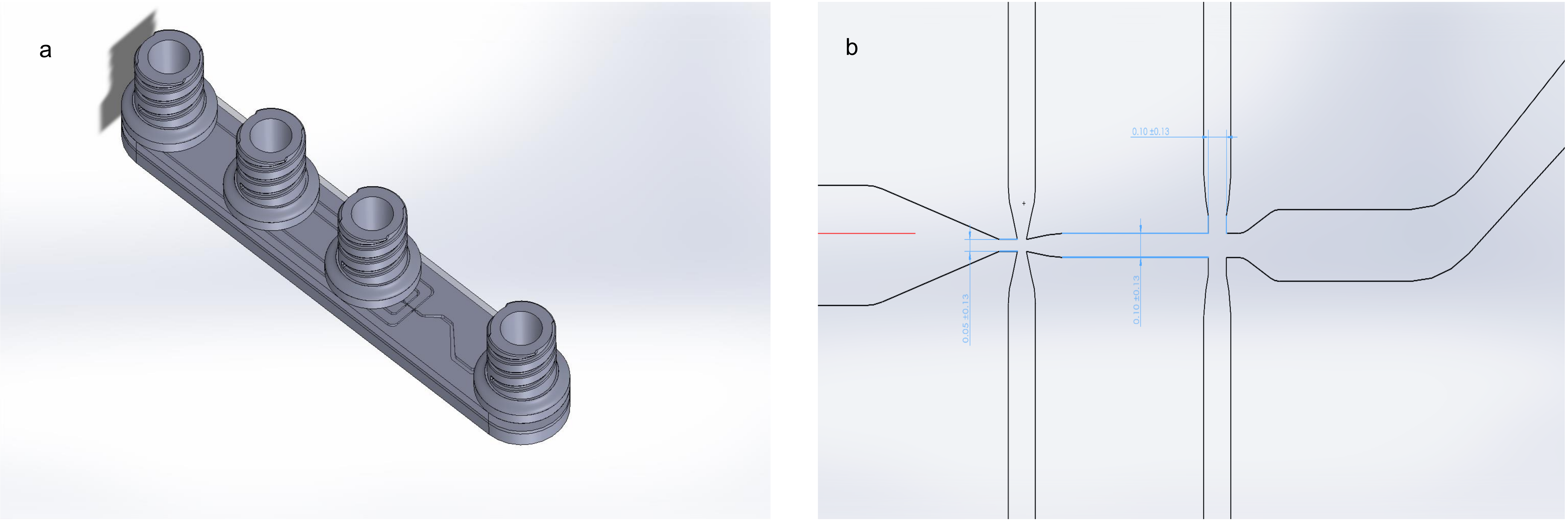
a
b
0.10 ±0.13
0.05 ±0.13
0.10 ±0.13

**Supplementary Fig. 1. Design of the dual flow-focusing microfluidic device used for nanoparticle synthesis.**

a) Two-dimensional schematic of the microfluidic channel layout illustrating the dual flow-focusing geometry and key channel dimensions used to generate W/O/W emulsions. The design enables controlled laminar focusing and rapid solvent exchange during nanoparticle formation.

b) Three-dimensional rendering of the microfluidic device showing inlet and outlet ports and overall device architecture.

The device design was adapted from previously reported flow-focusing microfluidic platforms for polymeric nano- and microparticle fabrication(Refs: 10.1007/s11051-021-05171-y; 10.1115/1.4026287; 10.1002/jor.26070).

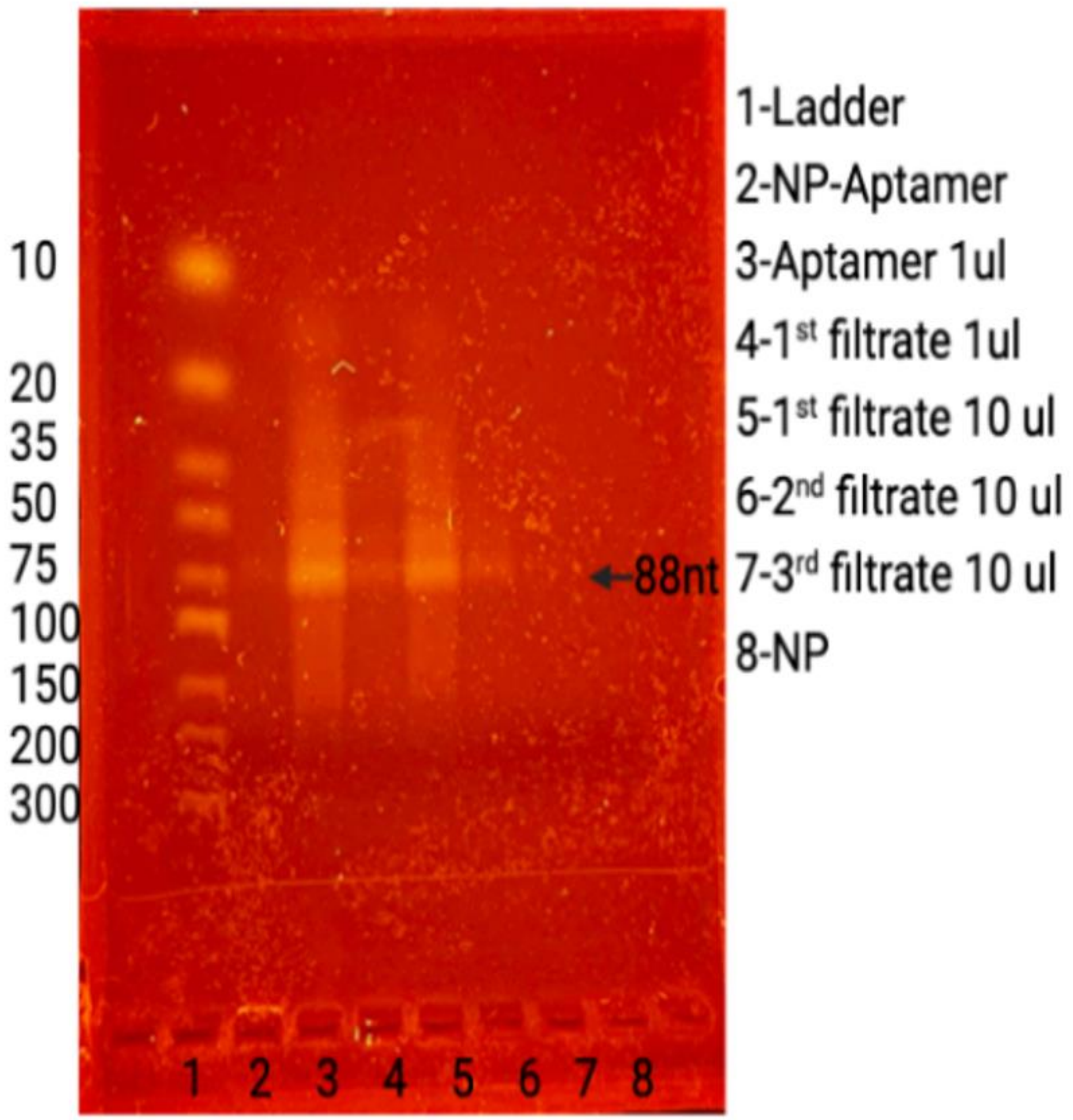


- **Supplementary Fig. 2. Gel electrophoresis confirmation of aptamer conjugation and washing efficiency.** Lane 1: DNA ladder; lane 2: aptamer-conjugated PLGA nanoparticles (mainly trapped in the well); lane 3: free aptamer; lanes 4–7: wash filtrates collected during purification; lane 8: unconjugated PLGA nanoparticles. Retention of aptamer-conjugated nanoparticles and reduced free aptamer signal in later wash filtrates support successful conjugation and removal of unbound aptamer.

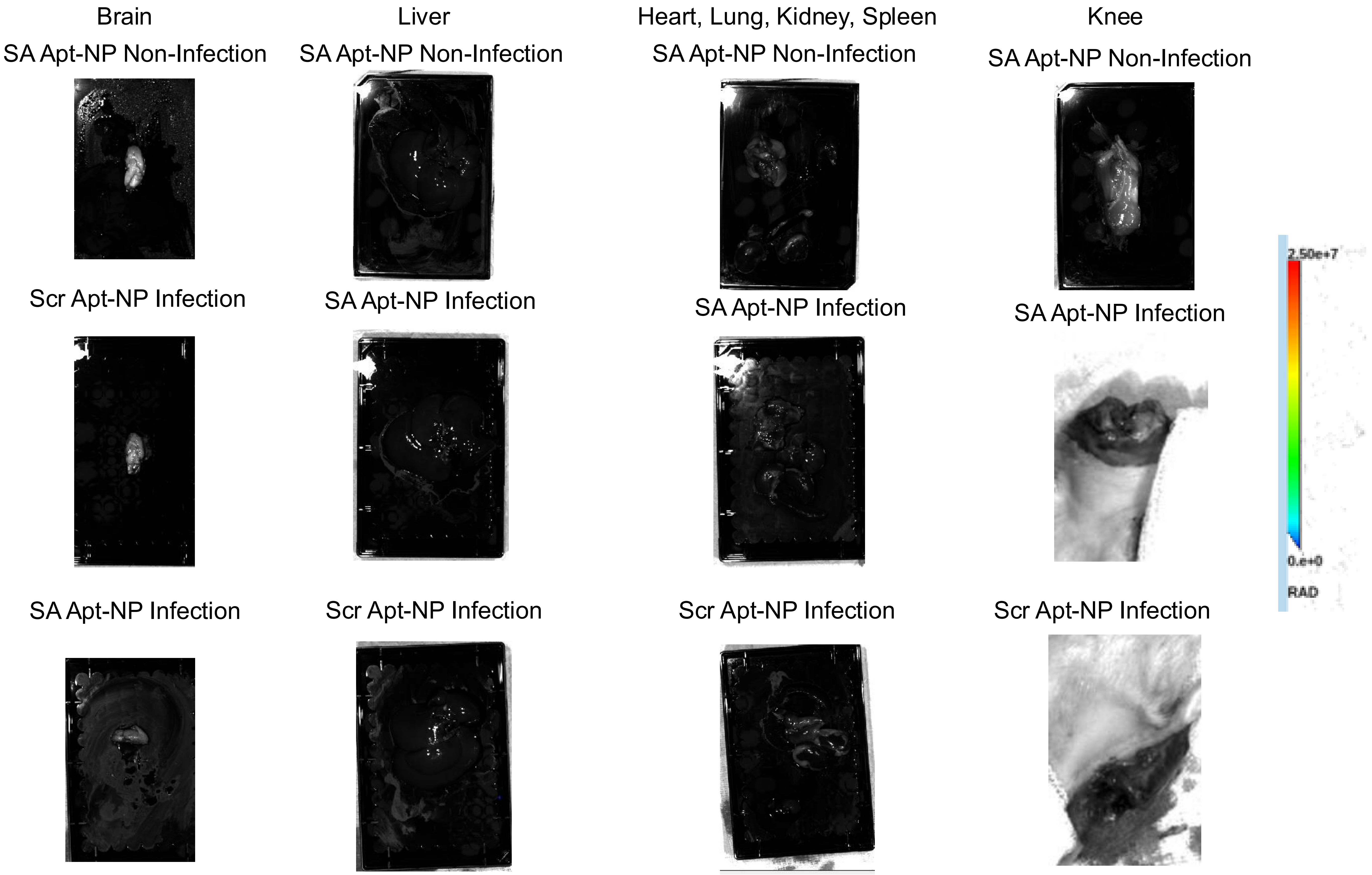
Brain
SA Apt-NP Non-Infection
Scr Apt-NP Infection
SA Apt-NP Infection
Liver
SA Apt-NP Non-Infection
SA Apt-NP Infection
Scr Apt-NP Infection
Heart, Lung, Kidney, Spleen
SA Apt-NP Non-Infection
SA Apt-NP Infection
Scr Apt-NP Infection
Knee
SA Apt-NP Non-Infection
SA Apt-NP Infection
Scr Apt-NP Infection
2.50e+7
0.e+0
RAD

**Supplementary Fig. 3. Representative ex vivo IVIS images of harvested tissues at endpoint.** Representative images of major organs and knee tissues collected after intra-articular administration of fluorescent SA Apt-NPs or fluorescent Scr Apt-NPs in infected and non-infected models. No appreciable fluorescence signal was observed in major off-target organs at endpoint; plate-reader quantification of extracted Cy5.5 from tissue homogenates was at or near background levels.

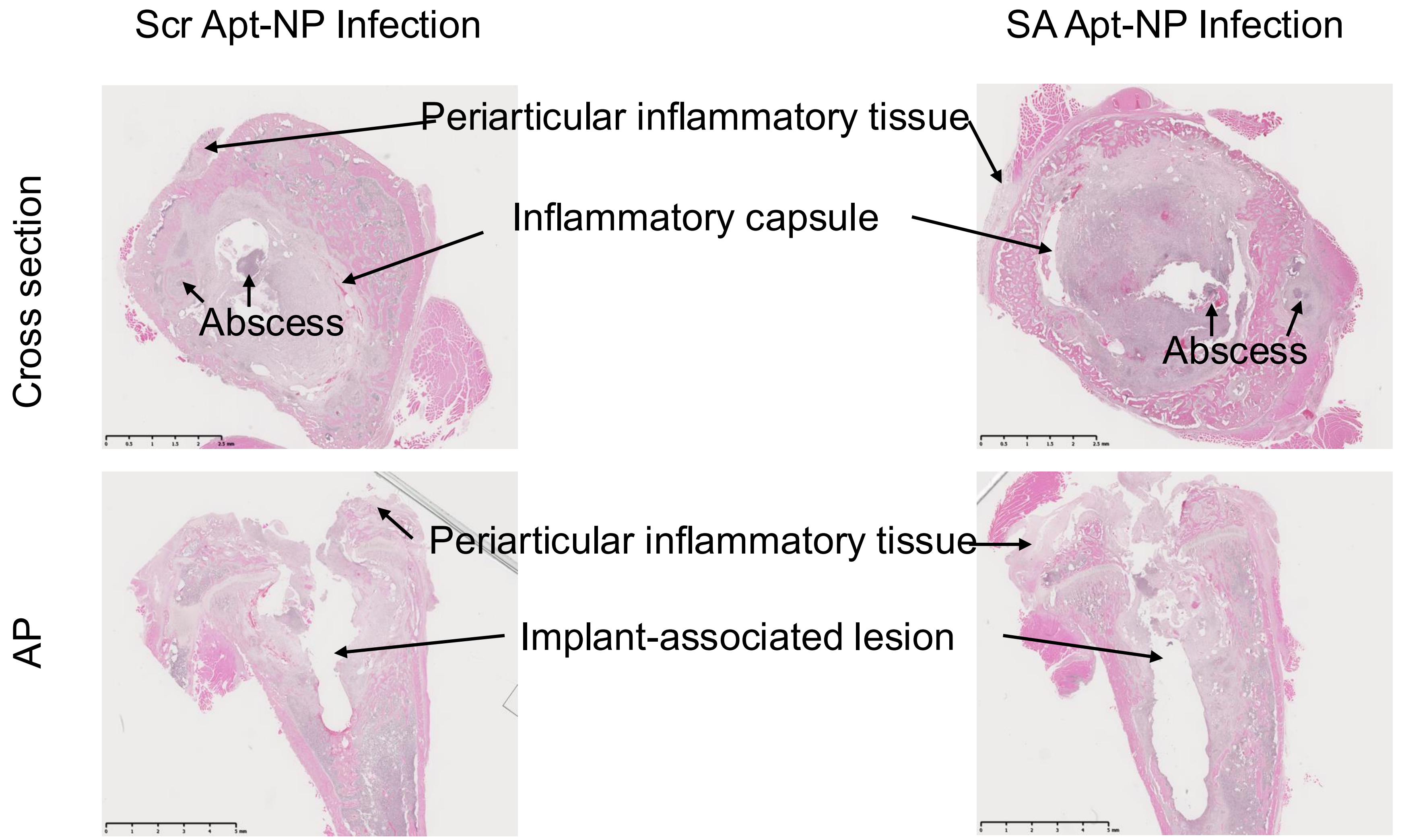


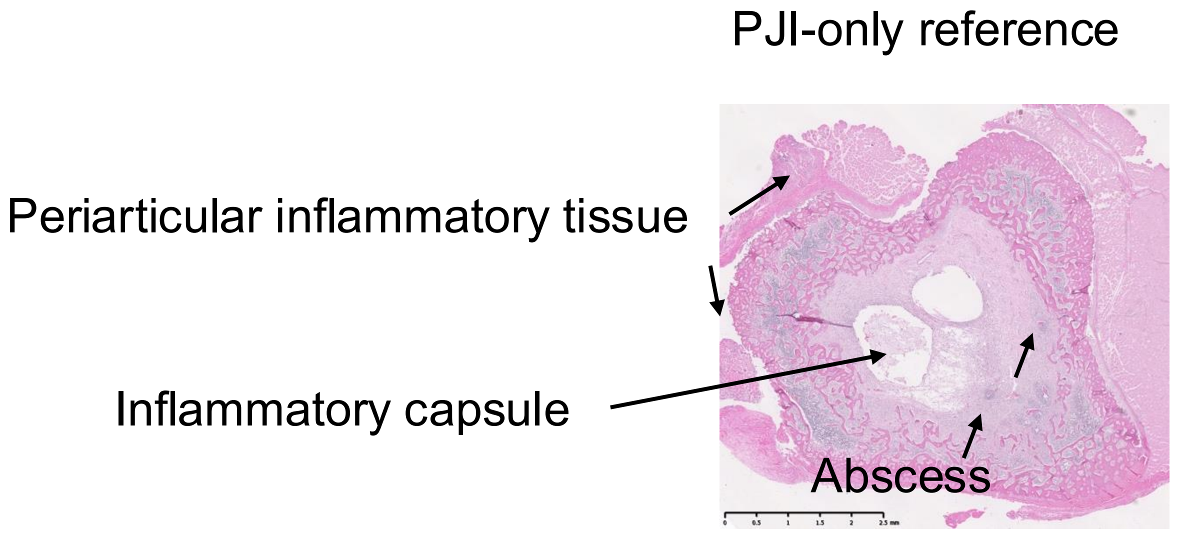


Reference only; not included in present experimental design

**Supplementary Fig. 4. Histologic evaluation of infected joints following Apt-NP injection with historical PJI reference histology.** Representative H&E-stained histologic sections from infected joints treated with scrambled Apt-NP or Staphylococcus aureus–targeted Apt-NP are shown in cross-sectional and anteroposterior (AP)-oriented views. Both Apt-NP–treated infected groups demonstrated periarticular inflammatory tissue formation and capsule-like remodeling, supporting successful establishment of the infection model. Qualitative histologic assessment did not show overtly increased inflammatory tissue expansion, suppurative change, necrosis, or structural tissue destruction in the S. aureus–targeted Apt-NP group compared with the scrambled Apt-NP group. A representative PJI-only histologic section from our previously published rat PJI model (Fan et al. 2020) is included as a historical reference to provide visual context for infection-associated histopathology. Arrows indicate infection-associated periarticular inflammatory tissue/capsule formation. Scale bars= 2.5 mm; 5mm.
